# Multi-compartment spatiotemporal metabolic modeling of the chicken gut guides the design of dietary interventions

**DOI:** 10.64898/2026.02.08.704450

**Authors:** Irina Utkina, Mohammadali Alizadeh, Shayan Sharif, John Parkinson

**Affiliations:** Program in Molecular Medicine, Hospital for Sick Children, Toronto, Ontario, Canada; Department of Pathobiology, Ontario Veterinary College, University of Guelph, Guelph, Ontario, Canada; Department of Biochemistry, University of Toronto, Toronto ON, Canada

**Keywords:** Microbiome, Metabolic Modeling, Poultry Gut, Dietary Interventions, Metabolic Interactions

## Abstract

Understanding the interactions between diet and the gut microbiome is critical for identifying dietary interventions that support gut health. This is of particular importance for poultry where the elimination of antibiotic growth promoters has resulted in an alarming rise in enteric infections with significant economic consequences. While the application of computational models capable of dissecting the metabolic interactions supporting gut communities has shown promise, they remain limited, largely ignoring the physiological and geographical considerations of the poultry gastrointestinal tract. To address these limitations, we developed the first multi-compartment, spatiotemporally-resolved metabolic model of the chicken gastrointestinal tract. Our six-compartment framework integrates avian-specific physiological features including bidirectional flow, feeding-fasting cycles, and compartment-specific environmental parameters. The model captured distinct metabolic specialization along the gut, with upper compartments enriched for biosynthetic pathways and lower compartments specialized for fermentation. *In silico* screening of 34 dietary supplements revealed context-dependent metabolic responses and predicted cellulose, starch, and L-threonine as robust enhancers of short-chain fatty acid production. A controlled feeding trial validated key predictions, particularly for butyrate, with prediction accuracy further improved through integration of trial-specific microbial community data. Our findings demonstrate that community composition is a major driver of metabolic outcomes and underscore the need for context-specific modeling. Our framework provides a mechanistic platform for rational dietary intervention design and is broadly adaptable to other animal or human gastrointestinal systems.

## INTRODUCTION

Over the past eight decades, antibiotic growth promoters (AGPs) have played an important role in livestock production, promoting gut health and reducing the burden of infectious disease^1^. However, the rise of antimicrobial resistance (AMR) associated with the use of AGPs (with AMR hotspots persisting across major poultry-producing regions in Asia, South America, and North America^2^) has created an urgent need for efficacious alternatives^3,45,62^. Research efforts have identified several promising alternatives, including phytochemicals, organic acids, probiotics, and prebiotics^3,4^. However, these interventions produce inconsistent outcomes across production systems, with substantial variability in efficacy between studies and farms^3–5^. This inconsistency reflects a critical challenge: intervention efficacy depends on baseline conditions, including diet composition, microbiota structure, and host physiology^5^. Consequently, identification of effective alternatives requires a more complete understanding of how potential interventions interact in the context with these additional factors.

At the core of these relationships are the metabolic interactions between gut microbiota, responsible for the degradation of dietary components and provision of nutrients, both to the microbial community as well as the host. Helping dissect these relationships are frameworks such as constraint-based metabolic models capable of explicitly simulating biochemical reaction networks. Recent advances have scaled these models from single species to community-scale frameworks capable of capturing interspecies interactions and metabolite exchange^6,7^. Tools such as MICOM and BacArena now enable quantitative simulations of microbiome metabolism under dietary perturbations^8–10^. In humans, these approaches have accurately predicted individual short-chain fatty acid (SCFA; metabolites associated with gut health^11^) profiles and revealed mechanistic links between microbial metabolism, host physiology, and diet^12,13^. Multi-compartment and agent-based frameworks have been established for mammalian systems, capturing pH gradients, oxygen diffusion, motility, and absorption along the gastrointestinal tract (GIT)^9,14–18^. Further, studies involving these platforms have shown that modeling the spatial organization of microbial communities improves predictive accuracy for metabolite fluxes and dietary intervention outcomes^14,16,17^.

Despite the limited application of community metabolic modeling to livestock, initial studies have shown much promise, revealing the impact of keystone taxa, as well as AGPs and their potential alternatives^19–21^. A significant limitation in these models, however, has been the focus on a single compartment, typically the ceca. Such models neglect the potential contributions of other compartments that feature distinct physical and chemical environments. Consequently, they do not capture the progressive processing of dietary components as they transit the GIT, or the functional contributions of individual taxa that may vary across sites – a limitation that is particularly relevant in the search for health-promoting probiotics that may colonize different regions^22,23^. The omission of these features impacts predictive accuracy for interventions affecting multiple gut sites and underscore the need for physiologically grounded, multi-compartment frameworks.

To address these limitations, we developed the first multi-compartment, spatiotemporally resolved metabolic model of the chicken GIT. Our six-compartment framework (gizzard, duodenum, jejunum, ileum, cecum, colon) integrates avian-specific physiological features, including bidirectional flow through peristalsis and reverse peristalsis, feeding-fasting cycles with diurnal shifts in gut motility, compartment-specific environmental parameters (pH, oxygen gradients, transit times), and nutrient absorption by the host. To reflect realistic variation in poultry production systems, the model was parameterized using 16S rRNA sequence survey data from chickens fed corn-or wheat-based diets, two nutritionally distinct formulations known to drive markedly different gut community structures and fermentative outputs^24–26^. The framework captured compartment-specific metabolic profiles, taxon-specific SCFA production, and temporal metabolite dynamics across feeding cycles. Systematic screening of 34 dietary supplements revealed context-dependent metabolic responses shaped by microbial community composition and gut region and identified cellulose, starch, and L-threonine as robust enhancers of SCFA production in the lower GIT. A controlled feeding trial provided direct experimental validation, with the accurate prediction of butyrate production. We further show that prediction accuracy increases with the use of community compositions representative of the birds being tested, that we suggest should be a prerequisite for developing precision nutrition strategies. Overall, our framework provides a mechanistic platform for rational design of dietary interventions to replace AGPs and advance strategies for sustainable poultry production.

## RESULTS

### Multi-compartment framework enables spatially resolved metabolic modeling

Previous metabolic models of chicken gut microbiomes have focused exclusively on cecal metabolism, neglecting how community composition varies across gut regions and how dietary nutrients are sequentially transformed during transit. To address these limitations, we developed a six-compartment model predicting both taxon-specific metabolic contributions and the fate of dietary inputs along the gastrointestinal tract. This framework connects adjacent gut sections, spanning from gizzard to colon, allowing the flow of both microbes and metabolites, and incorporates host physiological features including pH, oxygen gradients, motility, enzymatic digestion, and nutrient absorption.

Microbial community models for each compartment were built using 16S rRNA profiles from 10-day-old chickens fed either corn-(n=5) or wheat-based (n=5) diets. For each compartment, we selected representative taxa by identifying the most abundant bacterial families (comprising >1% relative abundance) and choosing the dominant taxa (represented as distinct Amplicon Sequence Variants (ASVs)) within those families that were mapped to high-quality chicken gut-derived MAGs^27–29^. Taxon selection was further informed by two additional criteria: documented prevalence in poultry microbiome literature and transcriptional activity in metatranscriptomic data from the same trial^5^. Genome-scale metabolic models for the resulting 29 bacterial taxa representing the core microbiota across compartments were reconstructed with gapseq^30^. To ensure the inclusion of key fiber-degrading pathways, models were additionally curated with carbohydrate-active enzymes derived from CAZyme predictions.

To account for physiological differences between gut sections, compartment-specific parameters such as pH, oxygen levels, and bacterial population sizes were set to reflect previously reported gradients along the tract (**Figure 1B**; see Methods). Nutritional inputs were calibrated to average daily feed intake and bacterial density, with feeding intervals matching poultry husbandry regimes (see Supplementary Methods). Transit of material (metabolites and taxa) between compartments was defined with reference to previous studies^31,32^. Nighttime dynamics, including reverse peristalsis, were simulated to mimic the fasted state and the reintroduction of digesta into the upper compartments. Finally, host-driven digestion and absorption of fatty acids, amino acids, and simple sugars were incorporated, with fractions removed from the relevant sections to reflect host uptake. Together, the developed comprehensive *in silico* framework provides a spatially and temporally resolved representation of microbial community dynamics in the chicken GIT.

**Figure 1.A.**
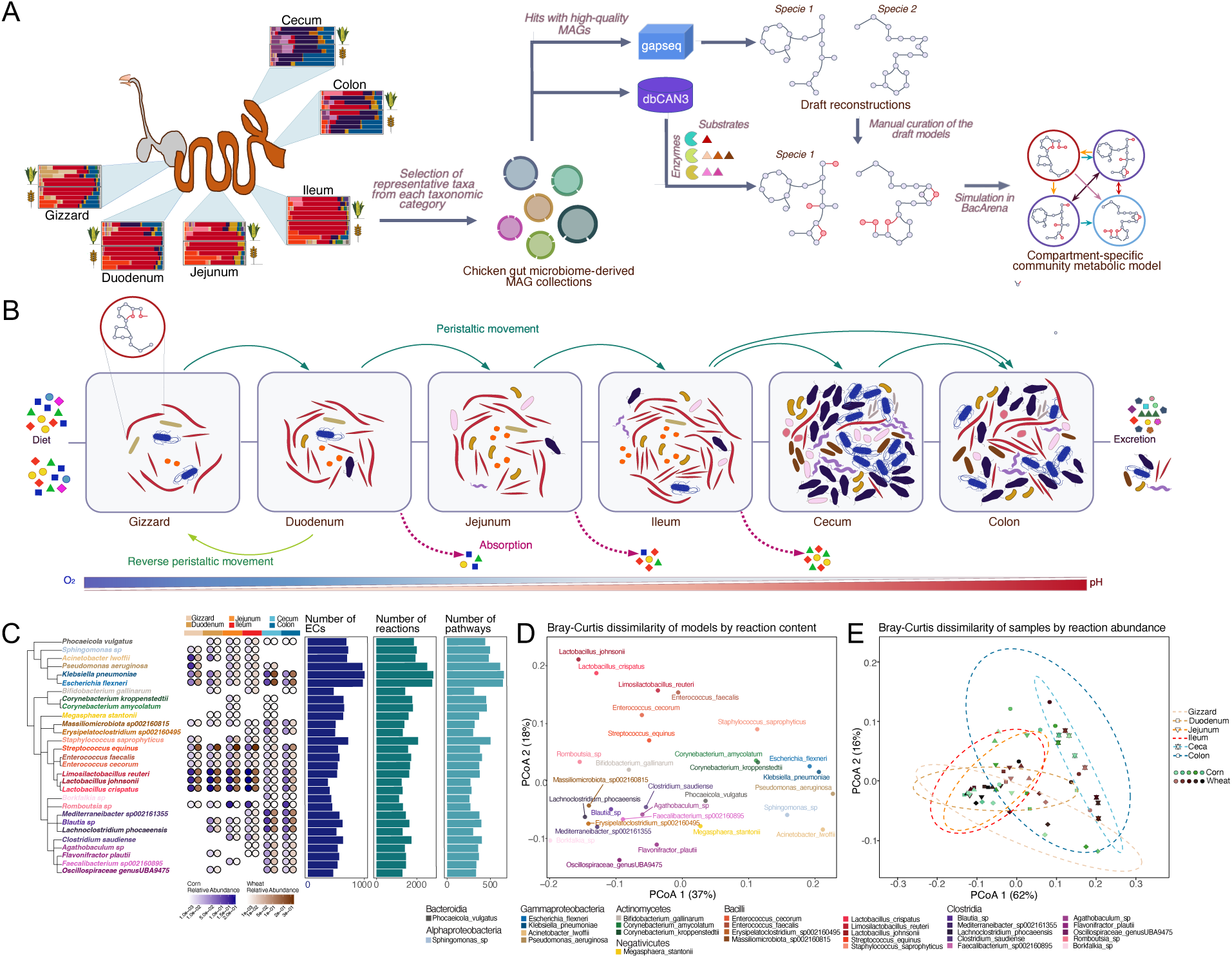
Overview of the modeling workflow. Community composition in each gut section (gizzard, duodenum, jejunum, ileum, cecum, colon) was parameterized using 16S rRNA profiles from corn-and wheat-fed chickens. Representative taxa were mapped to high-quality chicken gut-derived MAGs and reconstructed into genome-scale metabolic models with gapseq. Models were curated with CAZyme predictions (dbCAN3) to ensure representation of fiber-degrading pathways and integrated into sample-and compartment-specific community metabolic models. **B.** The six compartments are connected to simulate the flow of microbes and metabolites along the tract, with host physiological parameters including pH, oxygen, motility, digestion, and absorption incorporated to reflect site-specific conditions. **C.** Representative taxa selected to comprise core microbiota, shown in a phylogenetic tree with compartment-specific relative abundances (averaged across 10 samples) and the metabolic breadth of each corresponding GEM (number of EC numbers, reactions, pathways). **D.** PCoA of model reaction content using Bray-Curtis dissimilarity, illustrating clustering of taxa by their predicted metabolic repertoires. Each dot corresponds to a model of the taxon labeled. **E.** PCoA of reaction abundance profiles at the sample level, weighted by relative taxon abundance, showing clear separation of communities by gut compartment and partial separation by diet.

### Compartment-specific metabolic capabilities reveal distinct functional specialization along the chicken GIT

The 29 selected taxa span diverse phylogenetic groups and metabolic capabilities, with compartment-specific abundances reflecting the selective pressures of each gut region (**Figure 1C**). Metabolic capacities varied substantially across taxa: Enterobacteriaceae and other metabolic generalists (taxa with large, broadly distributed repertoires) encoded the highest numbers of reactions and pathways, while obligate anaerobes such as Oscillospiraceae displayed more specialized repertoires, reflecting adaptation to fiber fermentation in the lower gut. PCoA of enzyme content revealed clear metabolic stratification among the reconstructed models (**Figure 1D**). Metabolic generalists (*Escherichia, Klebsiella, Pseudomonas*) were separated from fermentation specialists (members of Lachnospiraceae and Oscillospiraceae families), with *Lactobacillus* species forming an isolated cluster likely reflecting their specialized fermentative metabolism. Notably, *Streptococcus* and *Romboutsia*, despite showing diet-dependent prevalence patterns in upper GIT compartments, clustered closely together, indicating metabolic similarity in the oxygenated, nutrient-rich upper gut. This suggests that taxonomic shifts driven by diet do not necessarily translate to functional divergence in the upper GIT communities. This contrasts with lower GIT communities, where taxonomic differences between corn- and wheat-fed birds (Lachnospiraceae/Clostridiaceae versus Enterobacteriaceae dominance) corresponded to distinct metabolic profiles.

Enzyme-level analysis further highlighted taxon-specific metabolic niches (**Figure S2**). Obligate anaerobes restricted to lower GIT compartments (members of Ruminococcaceae, Oscillospiraceae and Erysipelotrichaceae) encoded limited enzyme repertoires (510±62 ECs) concentrated in glycan metabolism pathways (331±44 pathways), consistent with specialization in cecal and colonic polysaccharide fermentation. By contrast, facultative anaerobes present throughout the GIT including cecum and colon (Enterobacteriaceae, *Pseudomonas*, and *Acinetobacter*) encoded substantially larger enzyme repertoires (913±145 ECs) spanning diverse metabolic pathways (600±81 pathways), reflecting their capacity as metabolic generalists. Their presence in the lower GIT likely supports obligate anaerobes through oxygen consumption and diverse metabolite production. Upper GIT-associated facultative anaerobes (*Streptococcus*, *Romboutsia*, *Lactobacillus*) encoded intermediate metabolic repertoires (505±114 ECs, 357±80 pathways), reflecting adaptation to rapidly available nutrients in the upper gut. Certain taxa, including *Sphingomonas*, *Acinetobacter*, and *Klebsiella*, uniquely possessed enzymes linked to xenobiotic degradation, suggesting ecological roles in processing dietary or environmental compounds that other members of the gut microbiome are unable to metabolize.

At the community level, PCoA based on reaction abundances revealed distinct clustering patterns that highlight metabolic specialization along the chicken gastrointestinal tract (**Figure 1E**). Although the metabolic profiles of the microbial communities differed significantly across gut compartments (PERMANOVA p=0.001), the pairwise differences between compartments were less pronounced (TukeyHSD p>0.05). The primary separation (PC1) distinguished upper from lower GIT communities based on contrasting metabolic strategies (**Figure S3A**). Upper GIT compartments were specifically enriched for bile salt hydrolases (glycocholate, glycochenodeoxycholate, taurodeoxycholate amidohydrolases), membrane lipid synthesis pathways (phosphatidate phosphatase, glycerol-3-phosphate cytidylyltransferase), and fatty acid biosynthesis (acetyl-CoA carboxylase), consistent with rapid biosynthetic activity in nutrient-rich, oxygen-replete conditions (**Figure S3B**). In contrast, cecal and colonic communities formed distinct clusters driven by enrichment in anaerobic fermentation pathways. Key metabolic specializations included SCFA synthesis (butyryl-CoA dehydrogenase, 3-hydroxybutanoyl-CoA oxidoreductase, butanoyl-CoA phosphotransferase for butyrate and propionate production), polysaccharide degradation (galacturonan hydrolases, arabinofuranosidases), and amino acid fermentation. The colon’s metabolic profiles partially overlapped with upper GIT profiles, reflecting the direct anatomical linkage between the ileum and colon, enabling the gradual transition in microbial activity and community composition as the digesta moves from the small to the large intestine (**Figure 1E**).

### Temporal dynamics of metabolite concentrations reveal compartment-specific signatures

Model simulations predicted taxon-specific SCFA production patterns that align with known metabolic capabilities (**Figure 2A**). *Escherichia flexneri* and *Klebsiella pneumoniae*, along with representatives of Lachnospiraceae and Erysipelotrichaceae families, were predicted to be among the major contributors to acetate production in the ileum, cecum, and colon. Acetate consumption by *Faecalibacterium* and *Clostridium* spp. fueled butyrate production^33,34^, while *Streptococci,* particularly abundant in wheat-fed birds, displayed metabolic flexibility, producing acetate in the more oxygenated ileal environment but consuming it under anaerobic, glucose-scarce conditions in the cecum and colon. Such shifts align with experimental evidence for environment-dependent regulation of Streptococcal acetate metabolism^35–38^. *Bacteroides* species emerged as the primary propionate producers in the lower GIT, corroborating the well-documented capacities of this genus^39^(**Figure 2A**). Together, these findings demonstrate that the six-compartment model captures both taxon-level metabolic repertoires and community-level metabolic specialization, including the adaptation of microbial metabolism to the varying physiological conditions of different gut sections and reliable prediction of SCFA production, consistent with previous reports.

**Figure 2.**
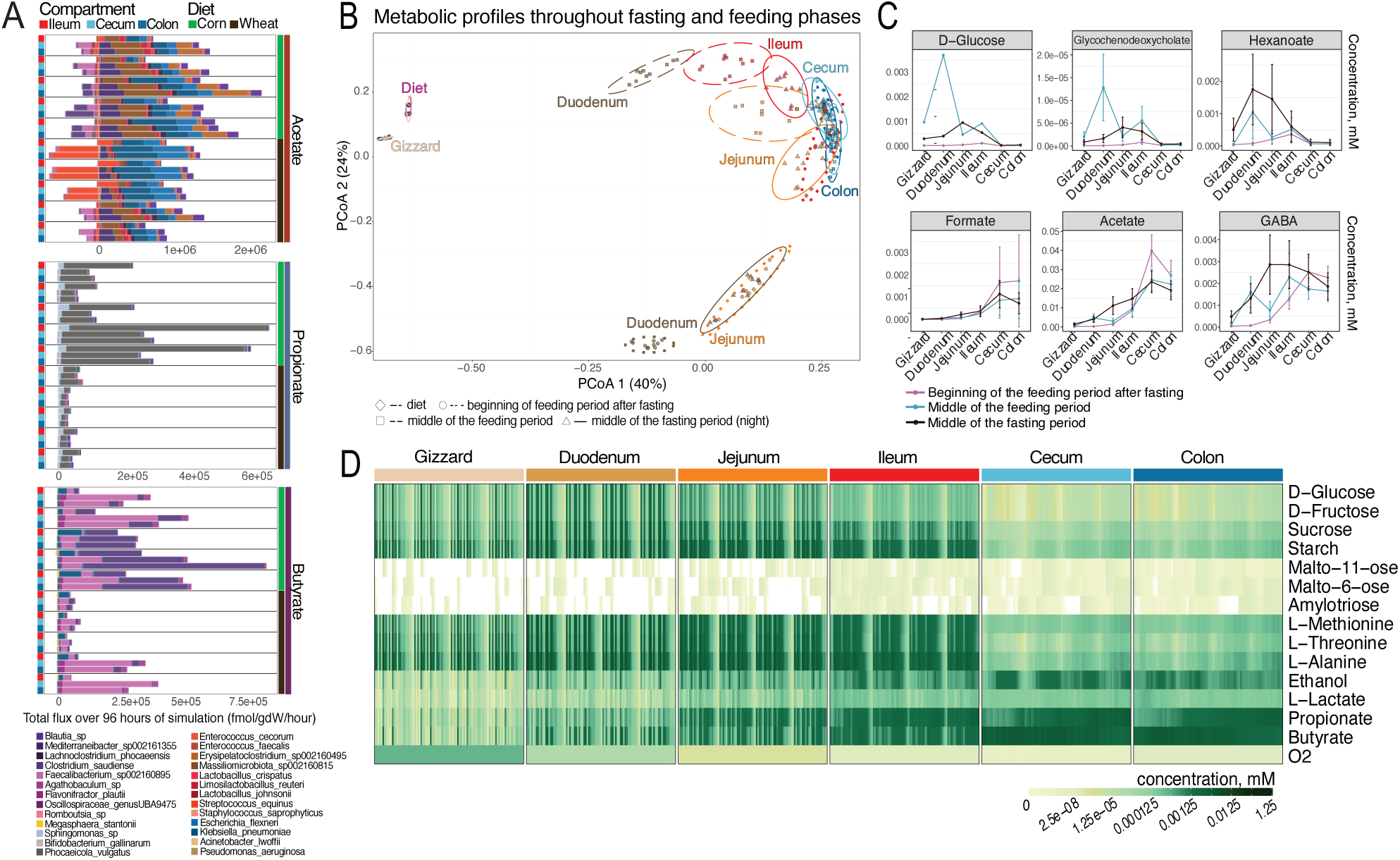
Temporal dynamics of metabolite concentrations across the chicken gastrointestinal tract during feeding and fasting cycles. **A.** Taxon-specific flux predictions for SCFAs over 96-hour simulations. **B.** Principal Coordinate Analysis (PCoA) of metabolite profiles across six gut compartments (gizzard, duodenum, jejunum, ileum, cecum, colon) at three time points: beginning of the feeding period after fasting, middle of the feeding period, and middle of the fasting period. Points are color-coded by compartment (corn-and wheat-fed samples combined to emphasize conserved compartment-specific signatures), with distinct clusters reflecting compartment-specific metabolic activities and shifts over time across both diets. **C.** Concentration profiles of selected metabolites across gut compartments at three time points: beginning of the feeding period (pink), middle of the feeding period (light blue), and middle of the fasting period (dark brown). The plots indicate the mean concentrations and standard deviations of metabolite concentrations from samples corresponding to chickens fed corn-based diet (n=5). **D.** Dynamics of metabolite concentrations across gut compartments during 96 hours of simulation for a selection of metabolites. Rows represent specific metabolites, with color intensity indicating concentration levels. The results shown were retrieved from simulation of one of the samples from corn-fed chicken.

Metabolite profiles across 96-hour simulations revealed dynamic compartment-specific signatures that shifted predictably with feeding-fasting cycles (**Figure 2B**). At the beginning of the feeding period, metabolite profiles clustered tightly within their respective compartments, reflecting the compartment-specific metabolic activity following fasting, with overlapping cecum and colon profiles, consistent with their similar environments and functionality. Notably, the unprocessed diet formed a distinct cluster, that remained separated from all gut compartments throughout simulations, demonstrating that microbial communities extensively transform dietary inputs into modified metabolite profiles rather than simply passing through dietary compounds unchanged. As feeding progressed, jejunal and ileal profiles showed increased dispersion, with the ileum shifting away from the cecum and colon and aligning more with the duodenum and jejunum, likely reflecting intensified nutrient processing as digesta transit accelerates. Cecal and colonic profiles remained stable and distinct, reinforcing their role as fermentation sites. During fasting, distal compartments maintained their clustering but shifted modestly toward the small intestine, while jejunal and ileal profiles converged, consistent with retrograde flow (reflux) of contents during periods of reduced peristaltic activity and nutrient scarcity, which redistributes metabolites backward along the GIT. The reflux mechanism is likewise responsible for the observed similarity in profiles between the duodenum and the gizzard.

Analysis of specific metabolite classes further illustrated these compartmental and temporal dynamics (**Figure 2C,D**). Simple sugars such as glucose, fructose, and sucrose accumulated in the duodenum and jejunum during feeding periods but were largely depleted in distal compartments. Residual glucose observed in the cecum and colon originated from the progressive breakdown of starch-derived oligosaccharides (maltoundecaose, maltohexaose, amylotriose). During fasting, glucose levels increased in the small intestine as a consequence of reflux and continued digestion of residual oligosaccharides. During the feeding period, amino acids including methionine, threonine, and alanine peaked in the duodenum and jejunum and declined downstream, reflecting rapid absorption in the small intestine; during fasting, reflux redistributed amino acids in upstream compartments, sustaining microbial growth in the absence of dietary inputs. Fermentation products (acetate, propionate, butyrate, ethanol, lactate and formate) reached the highest concentrations in the cecum and colon, particularly during fasting when the community relies on residual substrates from starch digestion and dietary fiber components.

Beyond accurately recapitulating concentration changes in primary substrates and fermentation products, the model also captured expected spatial and temporal dynamics of bile acids, fatty acids, and neuroactive metabolites. For example, concentrations of the secondary bile acid glycochenodeoxycholate peaked in the duodenum and jejunum during feeding and sharply declined in downstream compartments, while fasting caused modest increases in the upper compartments due to reflux (**Figure 2C**). Hexanoate concentrations peaked in the upper GIT (gizzard, duodenum, jejunum) during fasting, likely due to reflux-mediated redistribution, consistent with previous reports of higher medium-chain fatty acids in the chicken upper GIT relative to the lower GIT^40^. Interestingly, gamma-aminobutyric acid (GABA), a neurotransmitter produced by the gut microbiota and known for its neuroactive roles^41^, showed distinct temporal shifts across the GIT (**Figure 2C**). After fasting, GABA concentrations were highest in the cecum and colon (0.0025 ± 0.0007 and 0.0023 ± 0.0005 mM, respectively), reflecting sustained microbial production overnight. During feeding, concentrations increased in the duodenum (22-fold, to 0.0016 ± 0.0007 mM) and ileum (to 0.0023 ± 0.0011 mM). The jejunum showed the greatest fluctuation of GABA across the feeding-fasting cycle (Δ0.0025 mM), while cecal and colonic concentrations remained relatively stable. Such findings may have implications for the dynamics of GABA availability and its role in the gut-brain axis.

The time-dependent changes in metabolite concentrations observed *in silico* reveal how gut microbial communities respond to substrate availability and gut motility. Additionally, these temporal patterns highlight a critical methodological consideration: metabolite concentrations vary dramatically with sampling time. For instance, during fasting, SCFA levels peak in the cecum and colon, as fermentation intensifies and transit slows, allowing accumulation. Such findings underscore the importance of the timing of sample collection in experimental studies of gut metabolism^42^.

### Supplementation with L-threonine or starch is predicted to enhance SCFA production by diverse microbial communities

Establishing a model that recapitulates the spatial dynamics of metabolism in the chicken GIT, provides a valuable framework for examining the impact of dietary supplements. Of particular interest is understanding how differences in gut microbial communities may impact results and which supplements may be least susceptible to variations in these communities. With established roles in gut health^11^, we focused investigations on the production of SCFAs and screened 34 candidate supplements including established prebiotics (e.g., inulin^112^, XOS^113^, GOS^114^ and MOS^115^), non-digestible carbohydrates with documented beneficial effects on chicken gut health and performance (e.g., beta-glucans^116^, arabinooligosaccharides^117^, resistant starches^118,119^), essential nutrients commonly supplemented in poultry feed (e.g., thiamin^119^, riboflavin^120^ and zinc^121^), and metabolites identified through flux balance analysis as potential growth-limiting factors for beneficial bacteria (e.g., raffinose, L-leucine and pantothenic acid). Simulations involving supplementation of each metabolite were performed for 96-hours for each of the ten communities sampled (5 wheat-based and 5 corn-based with 5 technical *in silico* replicates performed for each community, see Methods).

Overall, we found that the metabolic response of each supplement varied according to both the supplement itself and the type of community (i.e., wheat-vs corn-based; **Figure 3A; Figure S4**). Further analysis of SCFA production fluxes demonstrated that multiple supplements could substantially alter SCFA production rates (**Figure 3B,C; Figure S4**). For instance, high doses of amino acid supplementation showed consistent results: L-threonine and glycine were predicted to significantly increase acetate production in both ileum and cecum irrespective of baseline community composition. Selected complex carbohydrates demonstrated similarly robust impacts: high doses of starch and cellulose were predicted to enhance production of all three SCFAs in both the ileum and cecum relative to controls. Several compounds displayed dose-dependent effects, with opposing outcomes at different concentrations (**Figure S4**). Most notably for propionate production: maltoheptaose supplementation at lower doses had minimal impact, while higher doses significantly decreased production, particularly in the cecum. Pantothenic acid showed similar concentration-dependent divergence, with low doses predicted to enhance SCFA production while higher doses reduced production. These opposing responses likely reflect shifts in community dynamics where higher substrate concentrations may favor fast-growing species that outcompete SCFA producers, reducing community-level yields. Conversely, certain commonly used additives in poultry production (e.g., riboflavin^43^, several mannooligosaccharides^44,45^) were unexpectedly predicted to decrease SCFA production across samples. Polysaccharide chain length also influenced SCFA production, with longer lengths of starch and mannooligosaccharides decreasing SCFA production. This is likely a consequence of longer chains requiring more specialized or complete sets of degradation enzymes, which are encoded by fewer bacterial genomes in the community.

**Figure 3.**
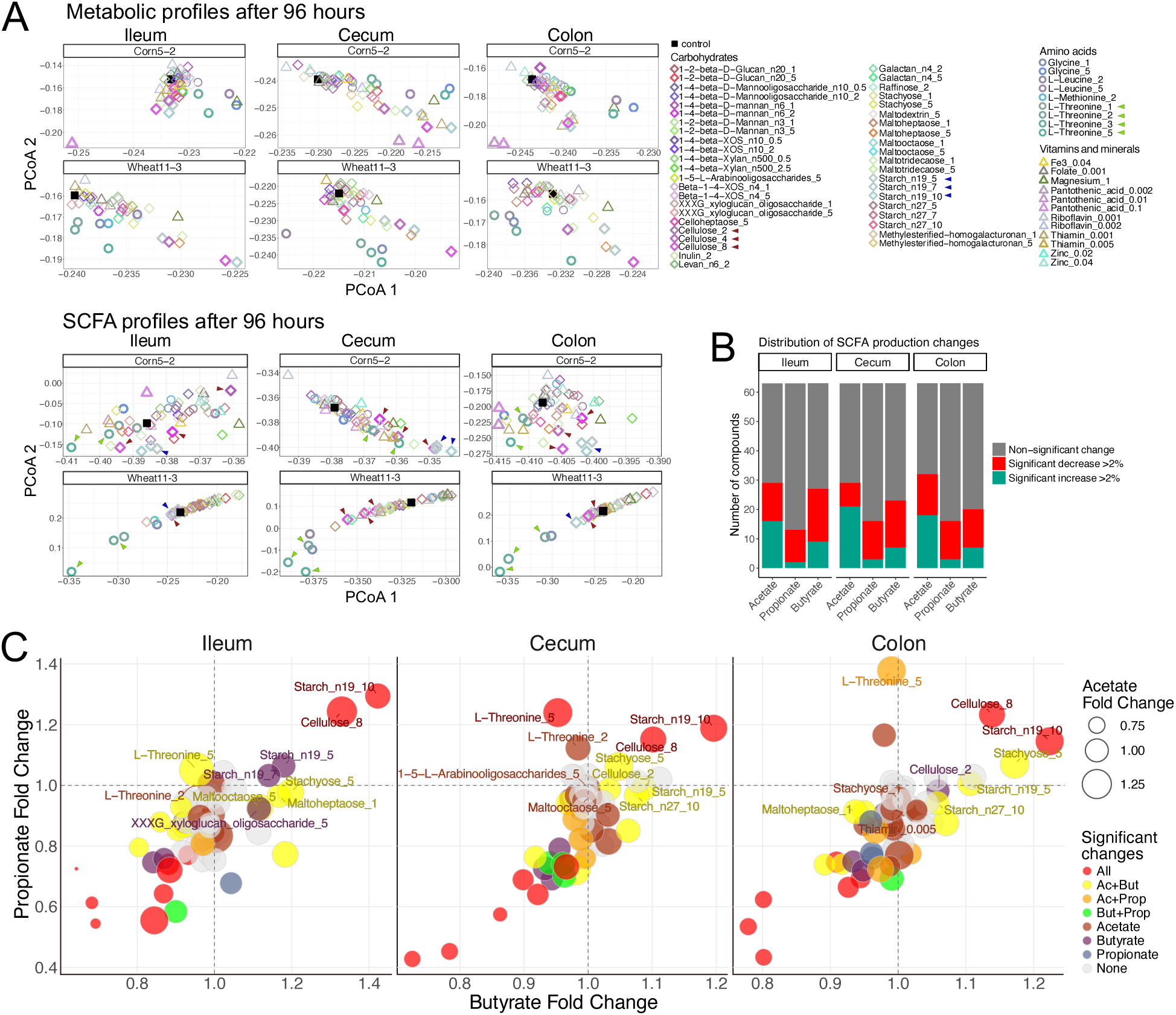
Impact of dietary supplementation on metabolite profiles and SCFA production across gut compartments. **A.** PCoA of metabolite concentrations after 96-hour simulations. Representative samples from corn-fed (Corn5-2) and wheat-fed (Wheat11-3) chickens are shown, comparing control conditions (black dots) with various dietary supplements across the ileum, cecum, and colon compartments. Key compounds discussed in the Results section (L-threonine, starch, pantothenic acid, cellulose, high-dose amino acids) are emphasized with thicker outlines. Arrows indicate L-threonine, cellulose, and starch - the three compounds selected for validation trials. Top panels: PCoA based on all metabolite concentrations. Bottom panels: PCoA based solely on SCFA concentrations. Supplement types are indicated by symbols and colors, with dosing shown after underscores (e.g., Inulin_2 = 2g per 100g diet). Complete dataset including all samples and compartments is presented in Supplementary Figure 4. **B.** Distribution of SCFA production responses to supplementation. Stacked bar plots show the proportion of tested compounds that significantly increased (green), decreased (red), or did not affect (gray) community-level SCFA production in each compartment. **C.** Scatter plots showing fold changes in butyrate (x-axis) versus propionate (y-axis) production for all tested supplements. Bubble size represents the magnitude of acetate fold change. Colors indicate which SCFA changes were statistically significant: red = all three SCFAs; yellow = acetate + butyrate; orange = acetate + propionate; green = butyrate + propionate; brown = acetate only; purple = butyrate only; blue-gray = propionate only; transparent gray = no significant changes. Dashed lines at fold change = 1.0 indicate average production levels in control simulations.

To identify promising candidates for potential application in the poultry industry, we focused on supplements that consistently increased SCFA production. Across compartments and diets, three supplements, namely, cellulose, starch, and L-threonine, were predicted to consistently enhance SCFA production (**Figures 3A,C; Figure S4B**). Cellulose, a complex polysaccharide, serves as a dietary fiber in poultry nutrition^46,47^ and is primarily fermented to produce SCFAs by cellulolytic bacteria such as Ruminococcaceae, Lachnospiraceae, and other Clostridiales^48^. Starch, a readily fermentable carbohydrate, is a major energy source for both the host and its gut microbiota. The digestion of starch starts in the small intestine, and a portion reaches the lower GIT, where bacterial enzymes degrade it into fermentable oligosaccharides and monosaccharides, subsequently converted to SCFAs through microbial fermentation pathways. L-threonine is an essential amino acid for chickens^124^. While primarily absorbed in the small intestine, a fraction of dietary L-threonine can reach the cecum and colon, where it serves as a substrate for bacterial species encoding threonine degradation pathways, yielding acetate and propionate as byproducts^49–51^. Based on their reproducible *in silico* effects across diverse microbial communities within the reference dataset, we selected these three candidates for validation in chicken feeding trials, testing two dietary doses of each to determine whether predicted metabolic benefits translated *in vivo*.

### Context-specific community models improved prediction accuracy

To evaluate the predictive capacity of our modeling framework, we performed a trial investigating the impact of two doses each of cellulose, starch, and L-threonine supplementation in broilers fed a corn-based diet. Ileal and cecal samples were collected at day 14 for 16S rRNA sequencing and targeted metabolomics (see Methods). Comparisons between predicted and experimentally derived metabolite concentrations revealed a spectrum of correlations (**Figure 4A**, upper panel). Among the metabolites tested, butyrate and fumarate concentration fold changes showed the highest agreement between predictions and experimental measurements (Spearman’s ρ=0.49 for both; **Figure 4B**). In contrast, acetate (Spearman’s ρ=-0.14), succinate (Spearman’s ρ=0.09), and α-ketoglutarate (Spearman’s ρ=-0.2) did not correlate with predictions, while lactate predictions were inversely correlated with trial measurements (Spearman’s ρ=-1.0, q=0.039). A potential confounder in these correlations is the microbial community composition, which differed between the validation trial (conducted in Guelph, Ontario) and the original trial used to construct the six-compartment models (conducted in Edmonton, Alberta). PCoA of cecal 16S rRNA profiles revealed distinct clustering of the trial-derived communities compared to both the original full-resolution dataset and the simplified composition used in the 6-compartment model (**Figure 4B**). PERMANOVA analysis confirmed statistically significant differences in community composition (p < 0.05) across both treatment groups and data sources, indicating substantial shifts in community structure between cohorts.

**Figure 4.**
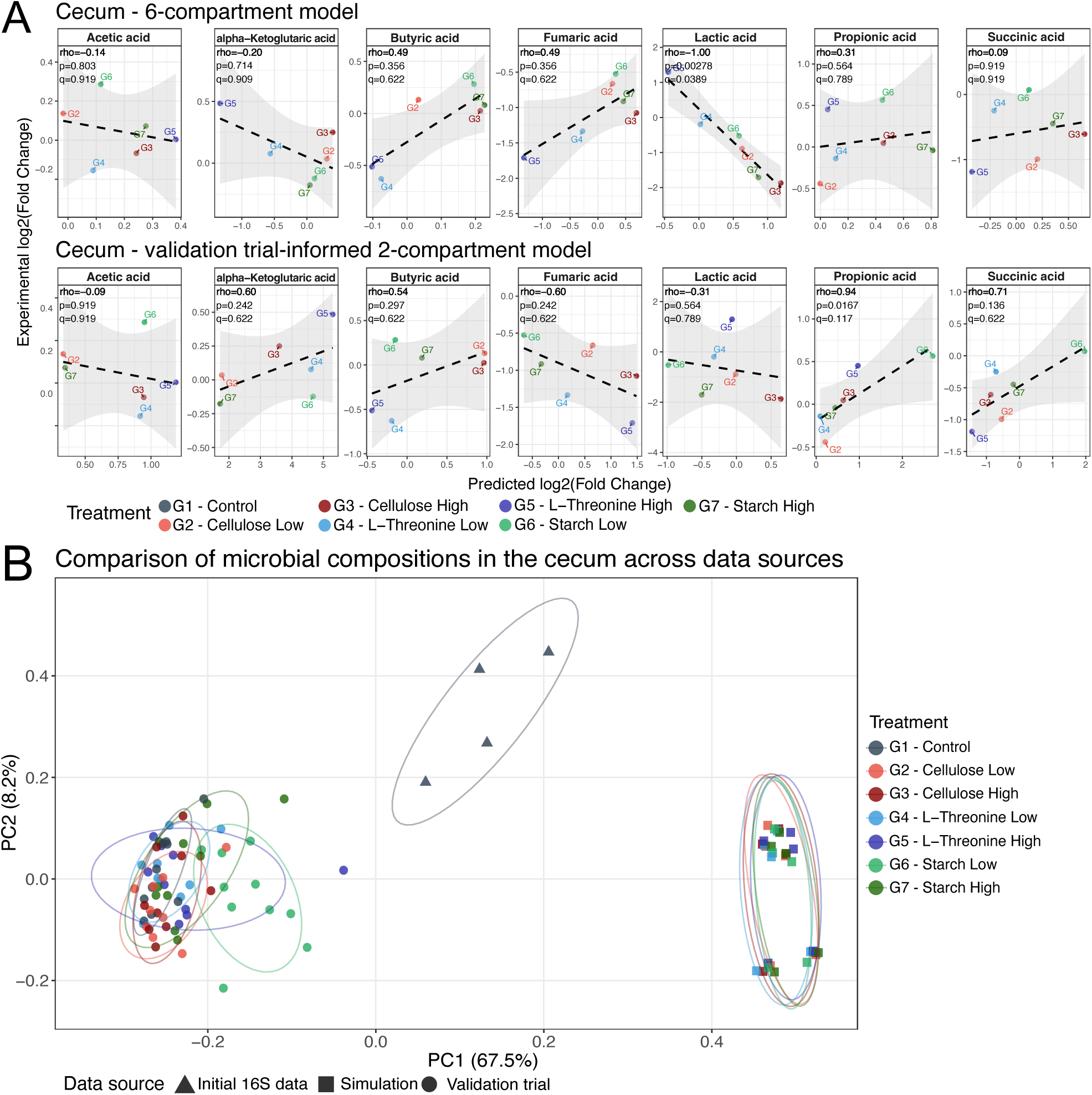
Trial-specific community composition data substantially improves metabolic predictions for cecal fermentation. **A.** Correlation between predicted and experimental log₂(fold change) values for seven fermentation metabolites in the cecum. Top row: Predictions from the 6-compartment model after 96-hour simulations compared to experimental measurements from the validation trial. Bottom row: Predictions from the trial-informed 2-compartment model after 48-hour simulations compared to experimental measurements. Each panel shows one metabolite with treatment groups indicated by color. Dashed lines represent linear regression fits. Spearman correlation coefficient (rho), uncorrected p-value (p), and FDR-adjusted q-value (q) are displayed for each metabolite. Fold changes were calculated relative to control group medians. **B**. Community composition differences. PCoA plots comparing taxonomic profiles between original reference data (Initial data), simplified model communities (Simulation), and trial communities (Validation trial) for cecum. PERMANOVA confirms significant separation between data sources (p < 0.05), highlighting taxonomic basis for model refinement.

These compositional differences reflected distinct taxonomic distributions between trials (**Figures S1, S6**). For example, Lactobacillaceae were more abundant in the original reference dataset (average 5.3 ± 3.9%, **Figure S1**) compared to trial communities (average 0.51 ± 1.88%, **Figure S6**) in cecal samples, contributing to predicted lactate accumulation in the original model and likely explaining the negative correlation for lactate. Notably, Enterobacteriaceae dominated the cecum in the reference dataset (average 21.3 ± 7.8%) but were nearly absent in trial communities (average 0.85 ± 1.8%). As facultative anaerobes, Enterobacteriaceae, mainly represented by *E. coli* in our models, produce acetate, succinate, and formate as major end products (**Figure 2A;Figure S8**) and are known to mediate cross-feeding networks in cecal communities^20^. The combined depletion of these families could alter both metabolite production and metabolic interactions, potentially explaining poor predictions for acetate and succinate. Conversely, Ruminococcaceae expanded from 4.2 ± 3.0% (reference) to 29.4 ± 8.4% (trial) in the cecal communities. This family comprises obligate anaerobic fiber degraders known to produce butyrate, including *Faecalibacterium*, a key butyrate producer in our models (**Figure 2A;Figure S8**). This may explain why butyrate predictions maintained reasonable correlation (Spearman’s ρ=0.49) despite the community restructuring. These findings demonstrated that while metabolic predictions were robust to variation within a reference cohort (**Figure 3**), community composition differences between facilities impacted prediction accuracy for certain metabolites, highlighting the need for context-specific modeling.

To address this limitation, we constructed a two-compartment model (reflecting the two compartments sampled in the validation trial – ileum and cecum) based on communities from the trial 16S profiles (see Methods). Trial-informed models were found to increase prediction accuracy across multiple metabolites (**Figure 4A**, lower panels). For instance, propionate correlation increased substantially from Spearman’s ρ=0.31 to ρ=0.94 (**Figure 4A**), aligning with elevated concentrations in the low-starch treatment group where *Bacteroides* were uniquely detected (**Figure S6)**. Similarly, predictions for succinate improved from Spearman’s ρ=0.09 to ρ=0.71, and α-ketoglutarate from Spearman’s ρ=-0.2 to ρ=0.6. Butyrate correlation showed modest improvement (Spearman’s ρ=0.49 to ρ=0.54), while lactate predictions shifted from significant inverse correlation (Spearman’s ρ=-1.0, q=0.03) to a more moderate mismatch (Spearman’s ρ=-0.31), consistent with reduced Lactobacillaceae abundance in trial-derived inputs. However, even with trial-specific community data, the models failed to accurately predict certain metabolites, revealing limitations beyond community composition effects. Acetate, which is broadly produced and absorbed throughout the gut, remained poorly predicted (Spearman’s ρ=-0.14 to ρ=-0.09), a pattern consistent with acetate prediction challenges reported in previous community modeling studies^52^. Fumarate correlation declined (Spearman’s ρ=0.49 to ρ=-0.60), with measured concentrations showing consistent depletion across all dietary treatments (ileum Kruskal-Wallis (KW) p=0.004, cecum KW p=0.002, significant post-hoc Dunn and Wilcoxon rank-sum tests, **Figure S7AB**), despite predicted net microbial production suggesting rapid microbial consumption or host absorption not captured in the model. In contrast, butyrate increases under low-dose cellulose and starch supplementation aligned well with predicted production by members of Ruminococcaceae, Butyricicoccaceae, and Lachnospiraceae families (cecum KW p=0.029), demonstrating accurate predictions when key producer taxa were present (**Figure S8**).

We note that high inter-individual variability and small sample sizes (n=6 per treatment) limited detection of statistically significant patterns for several metabolites, though trial-informed models consistently improved rank-order agreement with experimental trends. Together, these results demonstrate that community-matched metabolic models can reproduce *in vivo* fermentation patterns when grounded in appropriate taxonomic inputs, though host metabolic processes and biological variability remain important sources of prediction uncertainty.

## DISCUSSION

Optimizing poultry nutrition to promote gut health requires an understanding of how dietary inputs are transformed into metabolic outputs across the gastrointestinal tract. We developed a spatiotemporally resolved computational framework that simulates microbial community metabolism throughout the chicken GIT. This framework represents the most comprehensive spatial characterizations of gut metabolism in any livestock system performed to date. Our multi-compartment model captures three critical features: 1) spatial organization across the entire GIT through the simulation of six connected compartments; 2) temporal dynamics through inclusion of feeding-fasting cycles; and 3) the bidirectional flow of microbes and metabolites between neighbouring compartments through peristalsis and reverse peristalsis. Together these features represent a significant advance over previous metabolic modeling efforts in poultry that focused specifically on cecal metabolism^19,20^. By incorporating five additional compartments (gizzard, duodenum, jejunum, ileum, and colon), our framework treats the cecum as part of an interconnected system where upstream metabolic interconversions reshape substrate availability in the lower GIT and downstream metabolism. Our simulations revealed that metabolite landscapes and production rates vary substantially across compartments and over time – patterns that single-compartment models cannot capture. Upper gut compartments displayed distinct metabolic profiles enriched for bile salt hydrolases, membrane lipid synthesis, and fatty acid biosynthesis, while cecal and colonic communities specialized in SCFA synthesis pathways and polysaccharide degradation. The unprocessed diet remained distinct from metabolic profiles of all gut compartments throughout simulations, demonstrating that microbial communities extensively transform dietary inputs rather than simply passing compounds unchanged. Importantly, overnight reflux redistributes metabolites and helps explain convergences in metabolite profiles between adjacent regions.

Recent advances in mammalian gut modeling, including multi-compartment frameworks in mice^13,53^, human community-level FBA approaches^8,12,54^, and the CODY model of the human colon^16^, have demonstrated the value of spatially explicit and physiologically informed models for understanding host-microbiome metabolism. Our work builds on these concepts and extends them to poultry, where avian-specific physiological features, such as rapid upper GIT transit, steep oxygen gradients, and reverse peristalsis, play a defining role in nutrient processing and microbial ecology. Incorporating these traits allowed us to capture metabolite redistribution and temporal oscillations characteristic of the chicken gut. By modeling taxa individually rather than as a single metabolic entity, we predicted taxon-specific contributions to SCFA production: *Bacteroides* as the primary propionate producers, *Faecalibacterium* and *Clostridium* spp. as key butyrate producers, and Enterobacteriaceae and Lachnospiraceae as major acetate producers. Our study showed that the upper GIT (gizzard, duodenum and jejunum) communities were dominated by facultative anaerobes encoding fewer metabolic enzymes and enriched for bile salt hydrolases and fatty acid biosynthesis, reflecting adaptation to readily available nutrients under aerobic conditions. The ileum harbored mixed aerobic and anaerobic taxa, reflecting its transitional position between upper and lower GIT, while cecal and colonic communities were enriched in Clostridiales families with the capacity to ferment complex polysaccharides. Through the inclusion of spatial and temporal dynamics, our model captured metabolite accumulation associated with fasting and reflux. For example, fasting was associated with an increase in the production of SCFAs in the lower GIT (cecum and colon) as well as signaling molecules such as GABA in the upper GIT. These observations indicate that the timing of sample collection has the potential to substantially influence metabolite levels, consistent with recent studies showing that feeding cycles drive dramatic differences in metabolic measurements^42^.

Having established a spatiotemporally resolved framework, we leveraged its predictive capabilities to systematically screen dietary interventions. Simulations involving 34 candidate metabolite supplements revealed that their impacts were largely context-dependent, with metabolic responses varying by substrate type, concentration, polymer chain length, microbial community composition, and gut compartment. Some additives commonly used in poultry production (riboflavin, mannooligosaccharides) were unexpectedly predicted to decrease SCFA production, while others showed opposite dose-dependent effects (e.g., maltoheptaose, pantothenic acid). However, our model predicted three compounds – cellulose, starch, and L-threonine – to consistently enhance SCFA production across diets and compartments. The robustness of these predicted effects despite high inter-individual microbiome variation were tested in a poultry feed trial, enabling direct assessment of our model’s prediction accuracy.

Our experimental validation through a controlled feeding trial yielded critical insights into the determinants of predictive accuracy in community metabolic modeling. Butyrate responses to cellulose and starch supplementation aligned with model predictions, while other metabolites showed variable agreement with experimental measurements. For example, lactate accumulated in simulations but not *in vivo*, which reflected the much lower abundance of Lactobacillaceae in the trial cohort compared to the reference community used for model construction. Similarly, Enterobacteriaceae abundances were substantially lower in trial birds, while Ruminococcaceae increased 7-fold relative to the reference dataset, which could disrupt prediction for metabolites predominantly produced by specific taxa belonging to these families (e.g., acetate, butyrate). Since one of our main aims was to identify supplements that led to robust changes in SCFA production irrespective of community composition, this finding suggests that predictions may have limited relevance for poultry raised under different conditions and/or locales. By incorporating 16S rDNA survey data collected from the ileum and ceca during the trial within a more simplified two-compartment model, prediction accuracy improved, particularly for propionate, succinate, and α-ketoglutarate (Spearman’s ρ=0.31→0.94; ρ=0.09→0.71; and ρ=-0.20→0.60, respectively). This improvement underscores taxonomic structure as a primary determinant of metabolic function, aligning with recent observations that community composition drives differences in SCFA production^12^. At the same time, we note that even trial-informed model could not predict acetate (Spearman’s ρ=-0.09), consistent with challenges reported in other community modeling studies, or fumarate (Spearman’s ρ=-0.60), which showed consistent experimental depletion despite predicted microbial production, likely reflecting rapid host absorption not captured by our simplified fractional removal rates. This suggests future models could be improved through an improved understanding of host-metabolite absorption. These findings highlight that while metabolic predictions were robust to dietary and inter-individual variation within a reference cohort, cross-facility differences in community composition significantly impact accuracy, and host metabolic processes remain important sources of uncertainty.

Implementation-specific limitations are more readily addressable as data availability improves. The model’s reliance on automated genome-scale reconstructions means that enzymatic capabilities may be incomplete for underrepresented taxa. Our framework currently models only bacteria, excluding fungi and other microorganisms that may contribute to gut metabolism. The model assumes uniform distribution and mixing within the compartments, which may not accurately reflect the spatial microenvironments where localized metabolite gradients shape community interactions. Estimated rates of nutrient absorption and gut transit show inter-individual variation that our population-averaged parameters cannot capture. Additionally, the absence of host metabolic networks means we cannot represent how host processes (e.g., immune responses) affect microbial metabolism. Nevertheless, this modeling framework is adaptable and can readily accommodate future advances in microbial genomics and metabolic modeling. As new high-quality poultry MAGs and physiological measurements become available, they can be readily incorporated in future models. Incorporating gene and protein expression (metatranscriptomic and proteomic data) will further improve the accuracy of individual reconstructions, while coupling microbial models with host metabolic networks would improve representation of host absorption and host-microbiome metabolic exchange.

While our six-compartment model represents a significant step forward, it serves as a proof-of-concept for systematic modeling of the poultry GIT. By bridging *in silico* predictions with controlled feeding trials, we demonstrate both the promise and the limitations of current metabolic modeling approaches. This work lays the foundation for evaluating dietary interventions in poultry, guiding the design of targeted supplements that enhance SCFA production and gut health. More broadly, it illustrates how multi-compartment community models can accelerate hypothesis generation, reduce reliance on large-scale animal trials, and extend to other host-associated microbiomes where spatial structure and host physiology play decisive roles.

## METHODS

### Development of a six-compartment spatiotemporal model

#### Processing of 16S rRNA gene data

Reads were processed using the QIIME2^55^ package (version 2023.2), which involved trimming, filtering, and removing low-quality and chimeric sequences. The resulting high-quality data was then processed following the deblur pipeline. Sequences were clustered into amplicon sequence variant (ASV) groups and singleton sequences removed. Taxonomic classification was performed using the classify-hybrid-vsearch-sklearn function against the SILVA database (release 138.1)^55,56^. ASVs with an abundance less than 0.01% were removed to reduce the potential for observing bleed-through ASVs. ASVs identified as chloroplast or mitochondrial contaminants were also removed, and ASV tables were subsequently normalized and rarefied to a depth of 10,000 reads per sample to ensure comparability across datasets.

#### Selection of representative species and model reconstruction

To represent the most abundant taxa within each gut compartment, the top five ASVs per bacterial family were used as blastn queries against several collections of high-quality chicken gut-derived MAGs: a set of 12,339 MAGs from integrated 799 public chicken gut microbiome samples^28^, 469 MAGs from 24 chicken cecum samples^27^, and 461 MAG from five chicken gut compartments of 30 chickens^29^. Sequences meeting thresholds of 97% identity, e-value threshold of 1e-70, as well as passing assembly quality thresholds of ≥90% completeness, and ≤5% contamination were retained for further analysis. For ASVs that did not yield any MAG matches meeting these criteria, the 16S rRNA sequence was subjected to a blastn search against the NCBI RefSeq 16S database^57^. The highest scoring sequence alignments were then evaluated based on three criteria: (i) presence in chicken gut-derived MAG collections, (ii) prior documentation in poultry microbiome studies, and (iii) transcriptional activity confirmed by metatranscriptomic data from the same reference trial^5^. MAGs for taxa meeting these criteria and representing the most prevalent members of their genus in the chicken gut were selected for downstream metabolic model reconstruction. Complete details of genome sources, genome quality metrics, and taxonomic classifications for all selected MAGs are provided in Supplementary **Table S1**. Gapseq version 1.2^30^ with default settings was used to reconstruct genome-scale metabolic models. Draft metabolic models were curated manually to improve representation of carbohydrate metabolism (see Supplementary Methods).

#### Construction of sample-specific gut microbiota community models

Compartment-specific models were constructed for 60 samples (2 diets × 5 birds × 6 compartments) using the list of mapped taxa and their family-level abundances from 16S data. Representative species models were combined according to compartment-specific abundance profiles. In cases where sequencing data were unavailable for one compartment within a sample, the missing values were imputed using average relative abundances from the same dietary group and compartment. Samples missing data for more than one compartment were excluded.

#### Simulation framework and gastrointestinal configuration

All microbial community simulations were conducted using BacArena, an R-based platform for community modeling^9^. Bacteria were represented as individual agents occupying a 100 × 100 grid (10,000 cells total), where each cell could grow, move, and exchange metabolites with the surrounding environment. Growth optimization was performed for each individual at every time step using the microbial biomass reaction as an objective. Simulations were run for 96 hours (six-compartment model) or 48 hours (two-compartment model), each consisting of a 16-hour daytime phase (representing daylight exposure and feeding) followed by an eight-hour nighttime phase (mimicking sleeping and fasting conditions). To account for stochasticity in bacterial placement, movement, and replication, each bird-specific model was simulated with 5 replicates using different random seeds. The simulations were conducted with a time step of 1 hour, representing an hour of bacterial growth per iteration. To incorporate physiological factors, including absorption, peristalsis, reflux (reverse peristalsis), feeding, excretion, pH and oxygen homeostasis, the simulation environment was updated every hour (see more details in the **Supplementary Methods**). After each hour of simulation, these processes were executed to adjust the environment accordingly before proceeding with the simulation. The relevant parameters used during simulation initialization were defined for each compartment based on literature sources (**Table S2A).**

#### Comparative analysis of the functional capabilities of community models

To compare the metabolic capabilities across different gut compartments and samples, I first extracted the complete set of reactions from their respective community metabolic models. To quantify the metabolic potential within each sample, accounting for the varying contributions of different species, I summarized the reaction counts within each sample, where each reaction count was weighted by the relative abundance of the species contributing to that reaction. The resulting incidence matrices contained information on the reaction abundance in each community. Bray-Curtis dissimilarity matrices were constructed from the incidence matrices described using *vegdist* function from the vegan R-package, and Principal Coordinate Analysis was performed using *cmdscale* function from stats R-package.

#### Comparison of control simulations and simulations with supplementation

To evaluate the effects of different compounds on community fluxes and metabolite concentrations, fold changes were calculated for each combination of gut compartment, metabolite, and supplemented compound by dividing the metabolite concentration or community flux in the experimental group by the mean concentration or flux of 5 replicates from the control group. Additionally, given the experimental design, where validation is performed through chicken trials using 10 chickens per treatment, I subsampled 10 replicates from the original 25 by randomly selecting 2 out of 5 replicates from each of the 5 samples per diet. This subsampling approach resulting in 10 subsampled replicates per condition was designed to match the number of biological replicates to be used in the experimental validation and maintain the integrity of the statistical comparisons. Significant differences were determined by Wilcoxon signed-rank tests, that were conducted on the fold changes under the null hypothesis that the median fold change is equal to 1 (indicating no effect of the treatment). P-values were calculated using a two-sided test, and to account for the multiple comparisons inherent in this analysis, the Benjamini-Hochberg procedure was applied to control the False

Discovery Rate (FDR). Results were considered statistically significant at an FDR-adjusted p-value threshold of 0.05.

### Experimental validation and trial-informed model refinement

#### *In vivo* validation through chicken trials

While *in silico* simulations demonstrated significant effects of supplementation over 96 hours, the translation between *in silico* time units and actual community dynamics *in vivo* is not direct. Microbial growth rates, metabolic activities, and community interactions in the living gut are influenced by multiple physiological factors not fully captured in the model. Therefore, to ensure sufficient time for supplement-induced changes to manifest in the gut microbiome, a longer supplementation period was implemented in the validation experiments.

Seventy one-day-old broiler chickens were raised on standard corn-based starter diet for 7 days before random assignment to seven treatment groups (n=10 per group): control (G1), cellulose at 2g/100g (G2) and 8g/100g (G3), L-threonine at 1g/100g (G4) and 5g/100g (G5), and starch at 5g/100g (G6) and 10g/100g (G7). Supplementation lasted 7 days, birds were euthanized on day 14 and samples were collected from ileum and cecum, stored on dry ice, and then kept at −80 °C for microbiome analysis. All animal experiments were approved by the Animal Care Committee of the University of Guelph and adhered to the guidelines for the use of animals (Animal Utilization Protocol #4815).

#### DNA extraction

DNA was extracted from 10 mg of chicken gut samples (ceca and ileum) using the ZymoBIOMICS DNA miniprep kit with the addition of the ZR BashingBead (2.0 mm) tubes to increase homogenization and lysis of the samples. The extraction protocol was followed as indicated in the manual with a final elution volume of 50uL. 5 μL of the ZymoBIOMICS Spike-In Control I (D6320) was added to ileum samples and 10 μL to caeca. Positive (Zymo Microbial Community Standard) and negative (no input) extraction controls were run alongside the sample extractions.

#### 16S rRNA gene sequencing and processing

The V4 hypervariable region of the 16S rRNA gene was amplified using uniquely barcoded 515F (forward) and 806R (reverse) sequencing primers to allow for multiplexing^58^. Amplification reactions were performed using 12.5 μL of KAPA2G Robust HotStart ReadyMix (KAPA Biosystems), 1.5 μL of 10 μM forward and reverse primers, 7.5 μL of sterile water and 2 μL of DNA. The V4 region was amplified by cycling the reaction at 95**°**C for 3 minutes, 18x cycles of 95**°**C for 15 seconds, 50**°**C for 15 seconds and 72**°**C for 15 seconds, followed by a 5-minute 72**°**C extension. All amplification reactions were done in duplicate to reduce amplification bias, pooled, and checked on a 1% agarose TBE gel. Pooled duplicates were quantified using PicoGreen and combined by even concentrations. The library was then purified using Ampure XP beads and loaded on to the Illumina MiSeq for sequencing, according to manufacturer instructions (Illumina, San Diego, CA). Sequencing is performed using the V2 (150bp x 2) chemistry. A single-species (*Pseudomonas aeruginosa* DNA), a mock community (Zymo Microbial Community DNA Standard) and a template-free negative control were included in the sequencing runs. Sequencing data were processed using the identical QIIME2 pipeline detailed in section “Processing of 16S rRNA gene data” above, ensuring direct comparability between reference and trial datasets.

#### Targeted quantitative metabolomics data processing and analysis

A targeted quantitative metabolomics workflow was performed using a custom reverse-phase LC-MS/MS assay on an AB Sciex 5500 QTRAP® (Applied Biosystems/MDS Sciex) equipped with an Agilent 1290 UHPLC. Metabolites were identified and quantified by multiple reaction monitoring using isotope-labeled and other internal standards. The assay was prepared in a 96-deep-well plate format with an attached filter plate; the first 14 wells were allocated to one blank, three zero samples, seven calibration standards, and three quality control samples.

For all metabolites except organic acids, samples were thawed on ice and extracted. Extracts were loaded onto the filter plate, dried under nitrogen, derivatized with phenyl-isothiocyanate, incubated, and dried again. Metabolites were then extracted with 300 µL extraction solvent and collected by centrifugation into the lower deep-well plate, followed by dilution with MS running solvent. For organic acids, 150 µL ice-cold methanol and 10 µL isotope-labeled internal standard mix were added to extracted samples for overnight protein precipitation. Samples were centrifuged at 13,000×g for 20 min, and 50 µL supernatant was loaded into a 96-deep-well plate. 3-nitrophenylhydrazine reagent was added, followed by 2h incubation; BHT stabilizer and water were then added prior to LC-MS injection. Samples were analyzed by an LC method followed by a direct injection method. Data were acquired and processed in Analyst v1.6.3.

Dry-weight normalized concentrations were processed using MetaboAnalyst web platform^59^. Quality control filtering removed low-variance features (interquartile range < 5th percentile) and low-abundance features (median intensity < 5th percentile). The filtered dataset was normalized using probabilistic quotient normalization (PQN) with the control group (G1) as reference to account for dilution effects and technical variation. Fold changes for each metabolite were calculated relative to the control group median within each compartment. Statistical comparisons between treatment and control groups employed two-sample Wilcoxon tests with Benjamini-Hochberg FDR correction. To evaluate agreement between model predictions and experimental results, Spearman’s rank correlation coefficients were computed across all metabolites and treatment conditions, comparing predicted and observed fold changes within each compartment.

#### Construction of trial-informed 2-compartment models

From 16S rRNA data of 112 samples, 694 ASVs were filtered to retain those with ≥0.5% relative abundance, reducing sequencing noise. Functional profiles were predicted using PICRUSt2^60^, and ASVs were clustered by enzymatic similarity (Jaccard distance, 0.85 threshold), yielding 267 functionally distinct representatives. These were matched to high-quality MAGs through blastn-search against four chicken MAG catalogues: 461 MAG from five chicken gut compartments of 30 chickens^29^, MGnify chicken-gut v1.0 collection (13,386 genomes)^61^, a set of 12,339 MAGs from integrated 799 public chicken gut microbiome samples^28^, and 461 MAG from five chicken gut compartments of 30 chickens^29^. Matching was performed in three sequential passes on unmatched ASVs: (i) 97% sequence identity threshold with ≥80% query coverage; (ii) 95% identity with ≥70% coverage; (iii) 90% identity with ≥70% coverage. For each BLAST hit, a combined quality score was computed incorporating both sequence similarity (60% weight) and assembly quality (40% weight), where: sequence similarity = 0.4*identity + 0.3*coverage + 0.3*min[100,100×bitscore/500]), assembly quality = completeness - 5*contamination. Final selection included top-scoring hits per ASV, resulting in 170 unique MAGs. Complete details of genome sources, genome quality metrics, and taxonomic classifications for all selected MAGs are provided in Supplementary **Table S3**. Metabolic models were reconstructed using gapseq version 1.4.0^30^ with default settings for 44 bird pairs with sufficient data quality.

Bird-specific models paired ileal and cecal communities based on sample-derived ASV abundances, creating 88 compartment-specific models (44 birds × 2 compartments). Model assembly followed the same procedure as for the multi-compartment system, ensuring direct comparability.

### Definition of *in silico* dietary input

Nutritional information from the corn-based grower diet was adjusted to account for upstream digestion and absorption to create an *in silico* representation of dietary metabolites reaching the distal small intestine. The base diet composition was derived from the same poultry feed tables used in the 6-compartment model, with modifications to simulate the metabolic environment of the distal jejunum/proximal ileum. Metabolite concentrations entering the ileum were reduced based on published ileal digestibility coefficients for corn-based poultry diets. Mineral retention rates were set according to jejunal and ileal digestibility studies^62–68^. Similarly, carbohydrate retention reflected measurements on upper small intestine digestion^69–71^, while amino acids were retained at 25% based on typical ileal digestibility^69,72^. Fatty acid retention varied by saturation level^73^, and cellulose was retained at 50% to reflect partial fermentation in the upper gut. Essential metabolites (host-supplied metabolites, such as mucins, urea, and metabolites supporting growth of all community members) were added at baseline concentrations. The composition of the *in silico* diets used is available in the **Table S4**.

#### Design and simulation of a trial-informed 2-compartment (ileum-cecum) model

The design and configuration were identical to the six-compartment GIT simulation framework (described in the **Supplementary Methods**) with all the processes applicable to ileum and cecum implemented in the same way, except the reflux happening overnight. In the simplified 2-compartment model, reverse peristalsis was not considered due to the lack of upstream compartments and colon, involved in the process.

To maintain realistic metabolic growth rates, we implemented cooperative trade-off FBA (ctFBA) with L2 regularization, adapted from the MICOM framework^8^. This approach balances individual species growth rates against community-level biomass production through a two-step optimization. First, the maximum total community biomass production rate is determined, then the individual species growth rates are optimized while constraining total community growth to a fraction (τ, trade-off parameter) of the maximum, using L2 regularization to minimize growth rate variance. The trade-off parameter (τ) was calibrated independently for each compartment to ensure at least 90% of species achieved positive growth rates while maintaining physiologically realistic maximum growth rates (<3 h⁻¹ in the ileum, <5 h⁻¹ in the cecum). Calibration was performed at simulation initialization using the seeded microbial community and initial medium conditions, similar to MICOM’s approach.

The relevant parameters used during simulation initialization were defined for each compartment based on literature sources (**Table S2B**).

#### Simulation of dietary supplementation for validation experiments

To simulate dietary supplementation treatments, supplement concentrations entering the ileum were adjusted to account for upstream absorption and saturation effects. Cellulose supplementation at 2g/100g (G2) and 8g/100g (G3) was modeled as 1g/100g and 4g/100g reaching the ileum, maintaining an approximately 50% passage rate. L-Threonine supplementation at 1g/100g (G4) and 5g/100g (G5) was modeled as 0.4 g/100g and 2.5 g/100g reaching the ileum. While 40-60% of dietary threonine undergoes first-pass gut extraction, high doses saturate intestinal absorption, allowing greater proportions to reach the distal ileum. Starch supplementation at 5g/100g (G6) and 10g/100g (G7) in feed was modeled as 1.25g/100g and 2.0g/100g reaching the ileum, respectively, reflecting saturation of glucose absorption in the proximal small intestine at higher doses. These adjustments ensured physiologically realistic ileal concentrations while maintaining experimental dose-response relationships.

## DATA AVAILABILITY

For the 6-compartment model, 16S rDNA sequence datasets from a previous trial examining the impact of AGPs and diet on the poultry gut microbiome were downloaded from the SRA (project PRJNA614900). The previous study^5^ details the animal trial conditions, sequencing data retrieval, and 16S rDNA sequencing. The 16S rDNA sequence datasets generated in this study from the feeding trial are deposited under the BioProject PRJNA1403483.

## CODE AVAILABILITY

Scripts, source data and metabolic models required to reproduce the analyses of this study can be accessed through GitHub at https://github.com/ParkinsonLab/ChickenGIT_modeling

## ETHICS DECLARATIONS

### COMPETING INTERESTS

The authors declare that they have no competing interests.

## FUNDING

This work was funded by grants from the Natural Sciences and Engineering Research Council of Canada to J.P. (RGPIN-2019-06852/RGPIN-2025-06006), and Ontario Research Fund-Research Excellence to J.P. and S.S. High-performance computing was provided by the SciNet HPC Consortium; SciNet is funded by the following: the Canada Foundation for Innovation under the auspices of Digital Alliance Canada, the Government of Ontario, Ontario Research Fund–Research Excellence, and the University of Toronto. The funders had no role in the design of the study, collection of data and analysis, preparation of the manuscript, and decision to publish.

## AKNOWLEDGEMENTS

We thank the Centre for the Analysis of Genome Evolution & Function (CAGEF) at the University of Toronto for 16S rRNA amplicon sequencing services and The Metabolomics Innovation Centre (TMIC) for metabolomics profiling services.

## AUTHORS’ CONTRIBUTIONS

J.P. and I.U. conceived and designed the study. I.U., J.P., M.A. and S.S. designed the validation chicken trials; M.A. and S.S. conducted the chicken feeding trials, obtained regulatory approvals, and collected intestinal samples. I.U. performed metabolic model reconstruction and curation, simulations, data analyses, and visualization. J.P. and I.U wrote the manuscript. All authors reviewed and/or edited the manuscript.

## SUPPLEMENTAL MATERIAL

**Supplementary Figure 1. Taxonomic composition of gut microbiota in different gut compartments.** The relative abundance of bacterial families derived from 16S rRNA survey data is shown for samples from the gizzard, duodenum, jejunum, ileum, cecum, and colon, separated by corn and wheat diets. Color bars represent different taxonomic families, with relative proportions depicted for each gut compartment.

**Supplementary Figure 2. Distribution of enzyme commission numbers (ECs) across metabolic models.** The columns of the heatmap represent individual ECs, while the rows correspond to metabolic models representing different taxa. ECs are annotated using KEGG pathway designations, indicated in two layers to reflect primary and secondary pathway associations. For ECs lacking KEGG pathway annotation, MetaCyc pathway annotation was utilized when available. For ECs mapped to 3 or more major pathway categories, “Multiple superpathways” category was assigned. Each row represents a different taxon, annotated by taxonomic family.

**Supplementary Figure 3. Reaction-level drivers of compartmental separation observed in Figure 1E. A.** Distribution of PC1 (left) and PC2 (right) scores from the principal coordinates analysis (PCoA shown in **Figure 1E**) across gut compartments. PCoA was performed on Bray-Curtis dissimilarities of community metabolic reaction profiles weighted by species relative abundances. PC scores reflect the relative contribution of reaction sets associated with each axis rather than distances to compartment centroids. Dashed red lines indicate zero for each axis. PC1 primarily separates upper gastrointestinal tract compartments (gizzard, duodenum, jejunum, ileum) from lower gut compartments (cecum, colon), whereas PC2 captures additional metabolic stratification along the proximal-distal axis. **B.** Heatmap showing mean compartment-level abundances of metabolic reactions most strongly associated with PC1 and PC2 in Fig. 1E. Reactions were selected based on the highest positive and negative Spearman correlations between individual reaction abundances and PCoA axis scores (top loadings). Values are column-scaled (z-scores per reaction) to emphasize relative enrichment across compartments. Reactions associated with positive PC1 scores (upper GIT) include bile salt hydrolases, membrane lipid synthesis pathways, and fatty acid biosynthesis, consistent with biosynthetically active, oxygen-replete environments. In contrast, reactions associated with negative PC1 scores (lower GIT) are enriched for anaerobic fermentation processes, including short-chain fatty acid biosynthesis, polysaccharide degradation, and amino acid fermentation.

**Supplementary Figure 4. PCoA of metabolite concentrations after 96-hour simulations.** All 10 samples from corn-fed and wheat-fed chickens are shown, comparing control conditions (black dots) with various dietary supplements across all six gut compartments. Top panels: PCoA based on all metabolite concentrations. Bottom panels: PCoA based solely on SCFA concentrations. Supplement types are indicated by symbols and colors, with dosing shown after underscores (e.g., Inulin_2 = 2g per 100g diet).

**Supplementary Figure 5. SCFA production rates in simulations with supplementation.** Fold changes in total SCFA production by the microbial community over 96 hours of simulation when supplemented with a specific compound (y-axis), compared to the control, in ileum and cecum. The y-axis lists the various dietary supplements tested, with their concentrations indicated after the underscore (e.g., Inulin_2 denotes supplementation with 2g per 100g of diet). Each dot represents a fold change in community flux in a replicate (n=5) of a sample (n=10). Significance based on two-sided Wilcoxon signed-rank test with Benjamini–Hochberg correction for multiple testing: * – pvalue-adjusted < 0.05, ** – pvalue-adjusted < 0.01, *** – pvalue-adjusted < 0.001.

**Supplementary Figure 6. Taxonomic composition of gut microbiota in trial-derived samples from ileum and cecum gut compartments.** The relative abundances of bacterial families are derived from 16S rRNA survey data. Color bars represent different taxonomic families, with relative proportions depicted for each gut compartment.

**Supplementary Figure 7. Experimental validation of metabolite concentrations in response to dietary supplementation.** Targeted metabolomics measurements from chicken ileum and cecum samples following 7-day dietary supplementation. **A**. Box plots showing metabolite concentrations (μmol/g dry weight) across treatment groups for both ileum and cecum compartments. Each metabolite panel displays individual data points overlaid on box plots, with Kruskal-Wallis (KW) p-values shown in the top-left of each facet. Significant pairwise comparisons between treatment groups and control (determined by Dunn’s post-hoc test with Benjamini-Hochberg correction) are indicated by brackets with asterisks. **B**. Bar plots showing log2 fold changes relative to control group for each treatment in ileum and cecum. Asterisks indicate statistical significance from two-sample Wilcoxon rank-sum tests with Benjamini-Hochberg correction (* - p < 0.05, ** - p < 0.01, *** - p < 0.001). Fumarate showed consistent depletion across all treatments (KW p=0.002), contrasting with model predictions of net microbial production, suggesting rapid consumption or host absorption. Butyrate exhibited significant treatment effects (KW p=0.029), with increases under low-dose cellulose and starch aligning with predicted production by Ruminococcaceae, Butyricicoccaceae, and Lachnospiraceae. Lactate showed significant treatment effects (KW p=0.034) with high inter-individual variability, consistent with its role as both a fermentation product and rapidly consumed substrate. Succinate showed predominantly negative fold changes despite model predictions of increased production under starch (Bacteroidaceae) and cellulose (Lachnospiraceae), indicating additional metabolic sinks not captured in models. Acetate production increased in ileum, especially under cellulose and L-threonine treatments, however no consistent treatment effects were observed in cecum, reflecting acetate broad production across multiple families.

**Supplementary Figure 8. Predicted taxonomic contributions to metabolite production in the trial-informed 2-compartment model.** Bacterial flux predictions from 48-hour simulations of the trial-informed 2-compartment (ileum-cecum) model showing which taxonomic families contribute to production of seven fermentation metabolites. Horizontal stacked bar plots display cumulative metabolic flux (fmol/gdW/hour) for acetate, propionate, butyrate, lactate, fumarate, succinate, and alpha-ketoglutarate across all seven treatment groups (G1 (control), G2 (cellulose 2g/100g), G3 (cellulose 8g/100g), G4 (L-threonine 1g/100g), G5 (L-threonine 5g/100g), G6 (starch 5g/100g), and G7 (starch 10g/100g)) in ileum and cecum compartments. Each color represents a taxonomic family. Flux values represent total community production summed across all 48 hours of simulation and normalized by community size. The figure illustrates predicted treatment-specific modulation of bacterial metabolism, with cellulose and starch supplementation enhancing butyrate production capacity in cellulose-and starch-fermenting families, as referenced in the main text analyses.

**Supplementary Table S1. MAG selections for 29 bacterial species in the multi-compartment metabolic model.** Genome assignments for 29 bacterial species selected for genome-scale metabolic modeling of chicken gastrointestinal tract compartments. Species represent prevalent taxa from day 10 post-hatch samples across six GIT sites. Table includes species names, MAG identifiers, genome source databases (Intestinal Microbial Genomes ANI99 collection, HoloFood chicken gut MAG catalog), match quality categories, CheckM quality metrics (completeness, contamination), originating GIT site, assembly type, complete taxonomic classification, and selection notes. NCBI reference genomes were used when high-quality matches were unavailable in the three chicken gut MAG databases used as a reference.

**Supplementary Table S2. Parameters for spatial metabolic simulations using BacArena. A. Simulation parameters for the 6-compartment model.** Parameters for spatial metabolic simulations using BacArena. Table S2A provides parameters for the six-compartment gastrointestinal tract model representing gizzard, duodenum, jejunum, ileum, cecum, and colon. Parameters include initial bacterial community sizes per compartment, spatial grid dimensions (arena size), basal nutrient availability (essential metabolite concentrations), physicochemical conditions (pH, oxygen concentration), and dietary input schedules (proportion and timing of feed addition to gizzard). Additional parameters specify differential removal rates for day versus night periods and substrate-level removal percentages, reflecting circadian feeding patterns and intestinal motility in broiler chickens. **B. Simulation parameters for the ileum-cecum model.** Table S2B provides parameters for the simplified two-compartment ileum-cecum model, including bacterial removal rates and frequencies that simulate digesta flow and GIT transit. Additional parameters specify differential removal rates for day versus night periods and substrate-level removal percentages, reflecting circadian feeding patterns and intestinal motility in broiler chickens.

**Supplementary Table S3. MAG assignments for metabolic models in ileal and cecal microbial communities.** Representative MAG assignments for 267 functionally clustered ASVs from paired chicken ileum and cecum samples (validation trial). ASVs were clustered by 75% functional similarity (Jaccard index) of PICRUSt2-predicted enzyme profiles, with representatives selected to maximize MAG coverage. MAGs were selected from three chicken gut databases: ANI99 collection (Huang et al. 2022), MGnify chicken gut catalog (13,389 genomes), and HoloFood 12k MAG catalog. Selection prioritized combined BLAST quality (identity, bit score) and genome quality (completeness, contamination). Table includes: sample identifiers, treatment groups (control vs. prebiotic), body site (ileum/cecum), cumulative relative abundance per functional cluster, MAG identifiers, genome source databases, BLAST statistics (identity, bit score), MAG quality metrics (completeness, contamination), match quality categories (excellent/good/acceptable), and full taxonomic classifications.

**Supplementary Table S4. Diet compositions for metabolic simulations.** Complete metabolic compositions of corn-based and wheat-based grower diets used in simulations. Feed compositions from the original trial were translated into corresponding metabolites using animal feed tables mapping nutrient content to feed items. Oil compositions were sourced from vmh.life. Sugar and fiber categories were resolved into specific carbohydrates and polysaccharides based on published compositional studies, with cellulose used as the primary fiber source.

**Supplementary Table S5. Reactions added to and removed from metabolic models based on CAZyme gene annotations.** Manual curation of metabolic reactions to improve representation of complex carbohydrate metabolism beyond gapseq’s default modelSEED database. Sheet 1 documents 55 reactions added based on CAZymes detected by dbCAN3 analysis. Reactions were incorporated directly from modelSEED when exact matches existed, or manually defined using multiple biochemical databases for enzymes not in modelSEED.

Includes polysaccharide degradation reactions (arabinan, xyloglucan, xylan, etc.) and associated transport reactions, with detailed information on reaction definitions, EC numbers, stoichiometry, reversibility, affected species models, and newly introduced metabolites with chemical formulas. Sheet 2 documents 14 reactions removed from models when both: (1) the corresponding CAZyme was not detected by dbCAN3 in the genome, and (2) no significant sequence similarity was found during gapseq reaction search. Removed reactions include incorrectly predicted fructan biosynthesis and starch degradation activities.

## Notes

### Competing Interest Statement

The authors have declared no competing interest.

### Summary of Updates

Minor changes in the title; supplemental tables descriptions; list of references; quality of main figures

