## Supplementary Methods for "Multi-compartment spatiotemporal metabolic modeling of the chicken gut guides the design of dietary interventions"

### **Model curation**

Draft metabolic models were curated to improve representation of carbohydrate metabolism. As gapseq’s default biochemistry database (built on the basis of the modelSEED database^1^) contains limited annotations for complex carbohydrate utilization, additional reactions were introduced based on CAZyme (carbohydrate-active enzyme) annotations identified using dbCAN3^2^, which utilizes the expert-curated CAZy database^3^ as its foundation.

For each genome, the detected CAZymes were mapped to known biochemical reactions. When exact matches were available in modelSEED, they were incorporated directly; for enzymes not represented in modelSEED, reactions were manually defined using data from multiple biochemical reference databases^4–10^. In total, 55 reactions, including transport reactions, were added to the models where the corresponding enzymes were detected. Additionally, in certain cases, specific reactions were excluded from several models if they did not meet the following criteria: 1) the corresponding CAZyme was not detected in the metagenome used for reconstruction, as identified by dbCAN3; and 2) no significant sequence similarity to reference enzyme sequences was found during the gapseq reaction search. These modifications improved the models’ ability to metabolize complex dietary polysaccharides and better reflect known metabolic diversity of chicken gut bacteria (see **Table S5**).

### Design and configuration of the six-compartment GIT simulation framework

#### Initial ecosystem configuration

The initial microbial community composition in each gastrointestinal section is configured by setting compartment-specific parameters such as pH, oxygen levels, transit times and population size^11–15^. Due to computational constraints, the population size is scaled relative to the densest compartments, the cecum and colon, which are set to 4,000 individuals each. This scaling approach accounts for the order of magnitude differences in bacterial density across the different sections. Although the ileum has approximately a 100-fold lower bacterial density compared to the cecum^16^, representing it with only 40 individuals (a direct 100-fold reduction) would insufficiently capture the complexity and interactions of its microbial community. Therefore, we represent the ileal community with 1,000 individuals – a compromise that reflects a significant decrease in density while ensuring computational feasibility and adequate community representation. Similarly, the duodenal and jejunal microbial communities are represented by 500 individuals each, while the gizzard community consists of 250 individuals. This scaling strategy, though not maintaining exact proportions, allowed the model to capture the relative hierarchy of microbial populations across the various gastrointestinal sections while remaining computationally feasible.

#### Definition of in silico nutritional input

The feed composition used in the original trial^17^ was translated into corresponding metabolites using animal feed tables which map nutrient content to various animal feed items^18^. For oils not covered in these tables, their composition was sourced from the vmh.life resource^19^. The ‘Sugar’ and ‘Fiber’ categories in the feed tables were further resolved into specific carbohydrates and polysaccharides, with precise compounds and their proportions manually gathered from published studies^20–24^. Due to limited information on fiber in the chicken diet, cellulose was utilized as the primary source of fiber according to available literature^25,26^. . The composition of the *in silico* diets used are provided in the **Table S4**. A minimal growth-supporting medium was identified through flux variability analysis across all taxa, after which compartment-specific media were sequentially initialized, with the gizzard serving as the entry point for dietary components. After 24-hour initialization, compartment-specific metabolite concentrations were used as starting conditions for all compartments except the gizzard, which received simulated dietary inputs.

In order to simulate feeding regimes of a commercial broiler flock, portions of the daily feed intake were added to the gizzard every 2 hours, which aligns with the two common feeding strategies employed: *ad libitum* feeding, where chickens consume smaller meals more frequently throughout the day^27^ and restricted access, where chickens store larger quantities of feed in the crop which is then released to the rest of the GI tract in smaller portions^27,28^. The concentration of the input medium solution was calculated based on the average daily feed intake of a 10-day old chicken (approximately 100g^29^), the average weight of gizzard digesta (approximately 8 grams^30^)and the reported bacterial density in the chicken gizzard (ranging from 10^3^ to 10^6^)^16,30,31^. The initial gizzard microbial community was set to 250 individuals, dietary components were added nine times over the course of one day. The fraction of each feed intake is calculated as follows:

$$fraction of feed intake=\frac{100g \left( daily intake \right) \times250 (individuals in the in silico gizzard)}{9 \left( times per day \right)\times8g \left( avg digesta weight \right) \times5\times{10}^{4} CFU/g (avg bacterial density)} \approx7\times{10}^{-3}$$

This calculated value provides an initial estimate of input metabolite concentrations. Through iterative testing of concentrations around this value (±20%), we identified the optimal concentration that maintained stable community compositions across gastrointestinal sections while preventing unrealistic overgrowth.

#### Flow of luminal contents

The six-compartment GIT model incorporates parameters specific to each gastrointestinal compartment that define the characteristics of intestinal motility, including the frequency and magnitude of the peristalsis and reflux, which are implemented by transferring fractions of metabolites and bacteria between compartments. The proportion of metabolites and bacteria transferred between compartments was defined based on variations in transit time observed experimentally. Gizzard and duodenum were shown to have the fastest passage of digesta (~5-30 mins)^32^, while contents passage through the jejunum was recorded to be approximately 1 hour (25-60mins)^32,33^, and longer transit times observed in the ileum, up to 2 hours^32,34^. Cecal contents experience the longest transit time, being released only 2-3 times per day^35,36^, a crucial feature of the cecum's role as a central hub for bacterial fermentation and SCFA production.

During the daytime, the model simulates only gastro-duodenal reflux, with digesta moving from the duodenum to the gizzard. In contrast, the nighttime dynamics involve longer transit times^37,38^, implemented as smaller fractions of bacteria and metabolites being transferred between compartments, and includes reverse peristalsis across multiple GIT sections. During fasting periods, birds exhibit increased frequency of refluxes throughout the GIT, which allows digesta to be reintroduced to upstream compartments^39^. Accordingly, during nighttime simulations, the model implements both forward and reverse transfer of bacteria and metabolites between all compartments, while feed intake and excretion are suspended to reflect the fasted state. Finally, the model incorporated oxygen tolerance constraints for bacterial survival to address potential limitations in the metabolic models. Specifically, strict anaerobes (oxygen-sensitive species) that entered the aerobic gizzard via reflux were removed, as were obligate aerobes from the predominantly anaerobic cecum. This adjustment ensured the model accurately reflects the ability of specific bacteria to only occupy compartments compatible with their oxygen requirements.

#### Digestion and absorption of nutrients

Previous studies have shown that most digestive processes and nutrient absorption occur in the small intestine (duodenum, jejunum and ileum)^40^. To improve the accuracy of the model's predictions, fractions of fatty acids, amino acids, and simple sugars were removed from the corresponding gut sections based on known estimates of both host absorption and microbial-host degradation^41–47^. Additionally, the model incorporated known rates of carbohydrate processing by both host enzymes and bacterial communities. For instance, studies have shown that starch digestion and glucose absorption primarily occur in the small intestine, with the majority of digestion happening in the duodenum and jejunum (65-85% of starch) and nearly complete absorption (97% of glucose) taking place by the terminal ileum^41,42,48,49^. The duodenum is the primary site of glucose absorption, which then continues at a decreased rate down the small intestine^41^. The metabolic rates were assumed to be consistent throughout the day and night^50^, and therefore, the absorption and digestion rates remained the same. Furthermore, the environment in each compartment was considered to be constantly mixed, allowing bacteria to move to any other position within the compartment. Metabolite concentrations were evenly redistributed after each hourly iteration, resulting in a uniform distribution of metabolites in each grid cell.
