## Supplementary figures and images for "Multi-compartment spatiotemporal metabolic modeling of the chicken gut guides the design of dietary interventions"

### Supplemental Figure 1

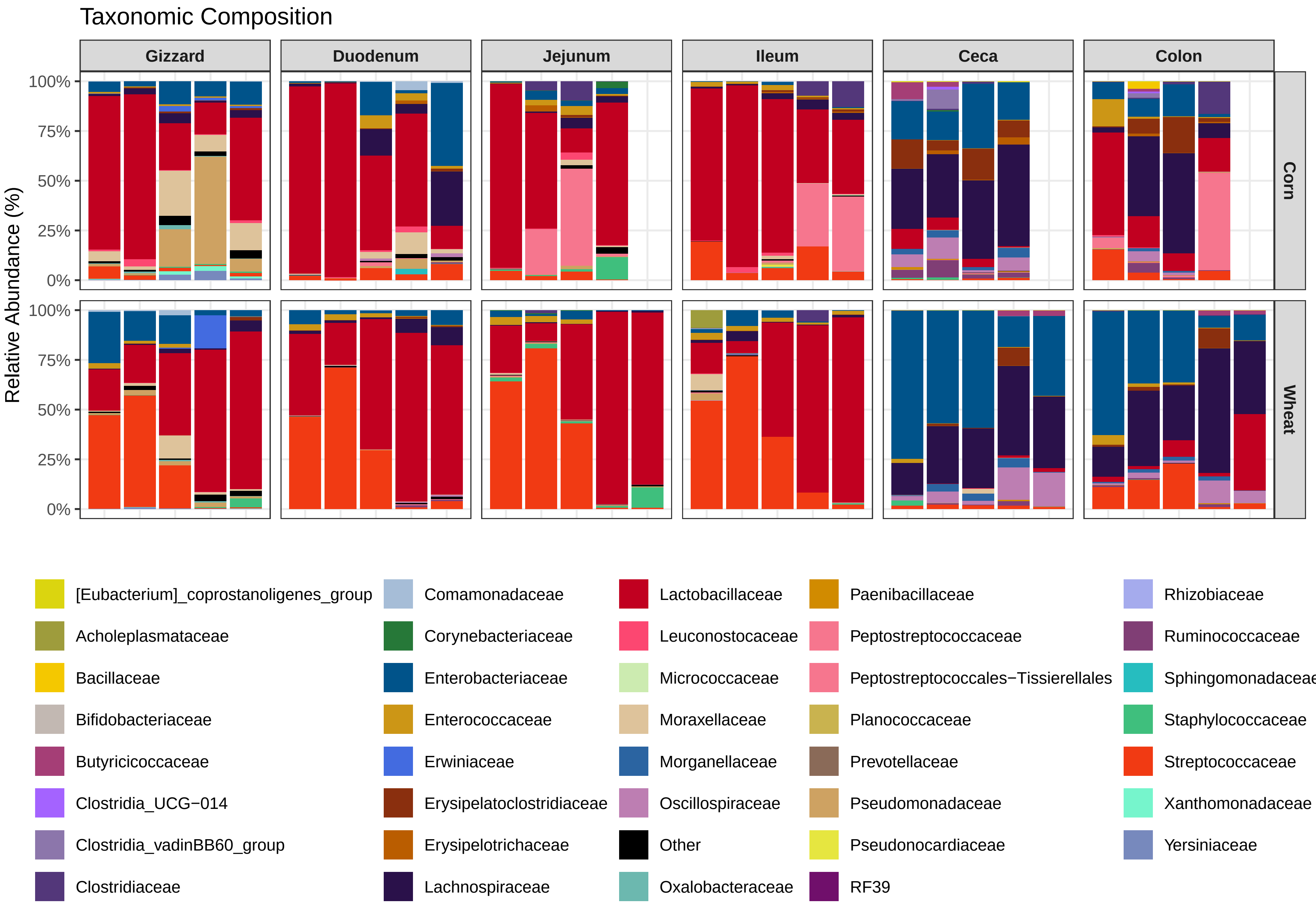

### Supplemental Figure 2

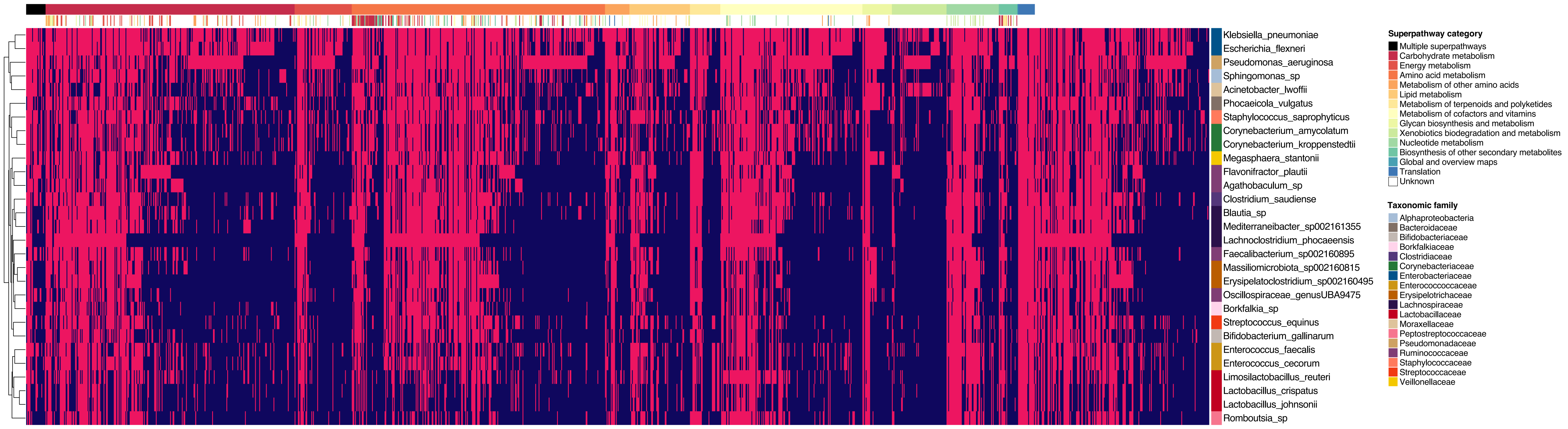

### Supplemental Figure 3

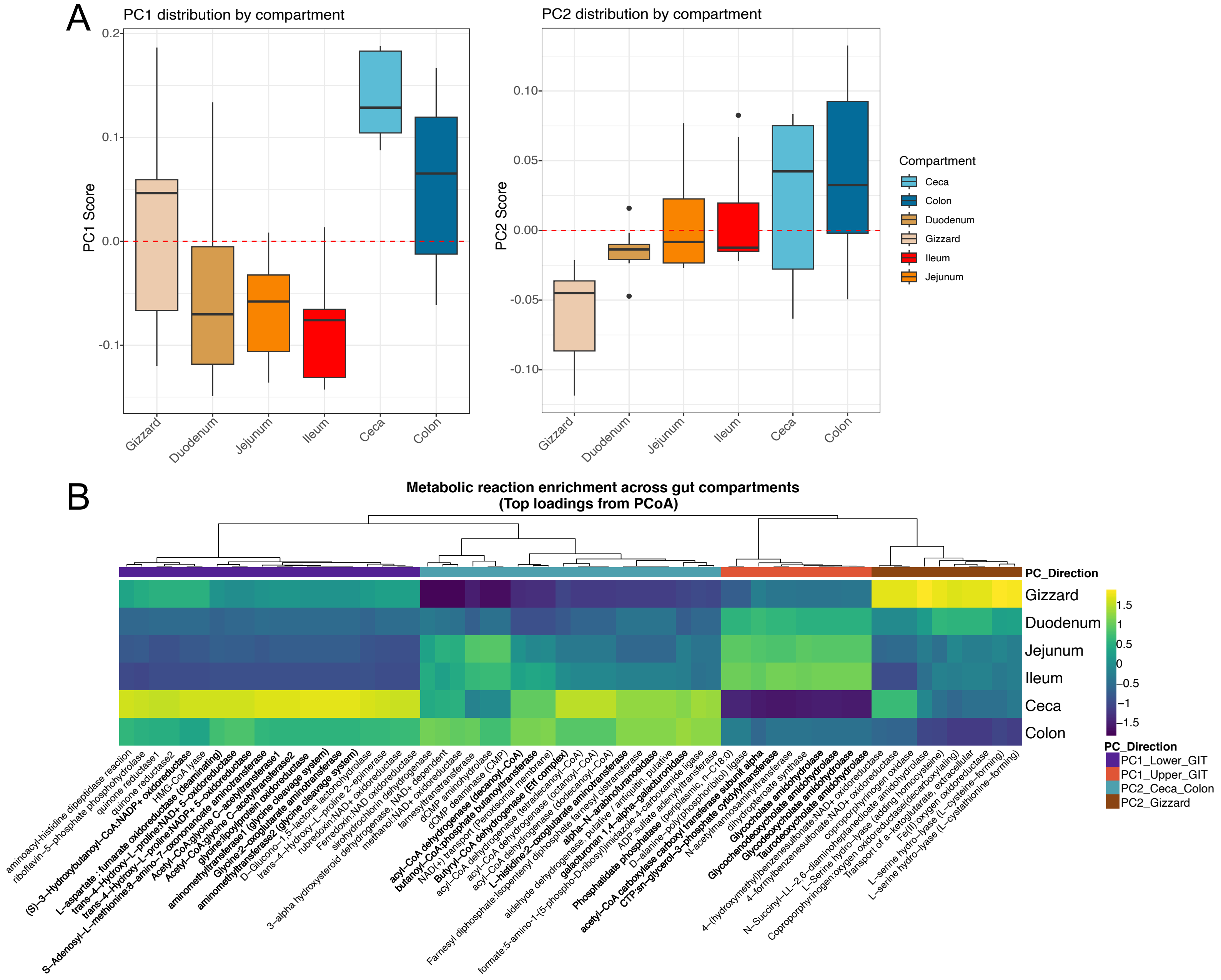

### Supplemental Figure 4

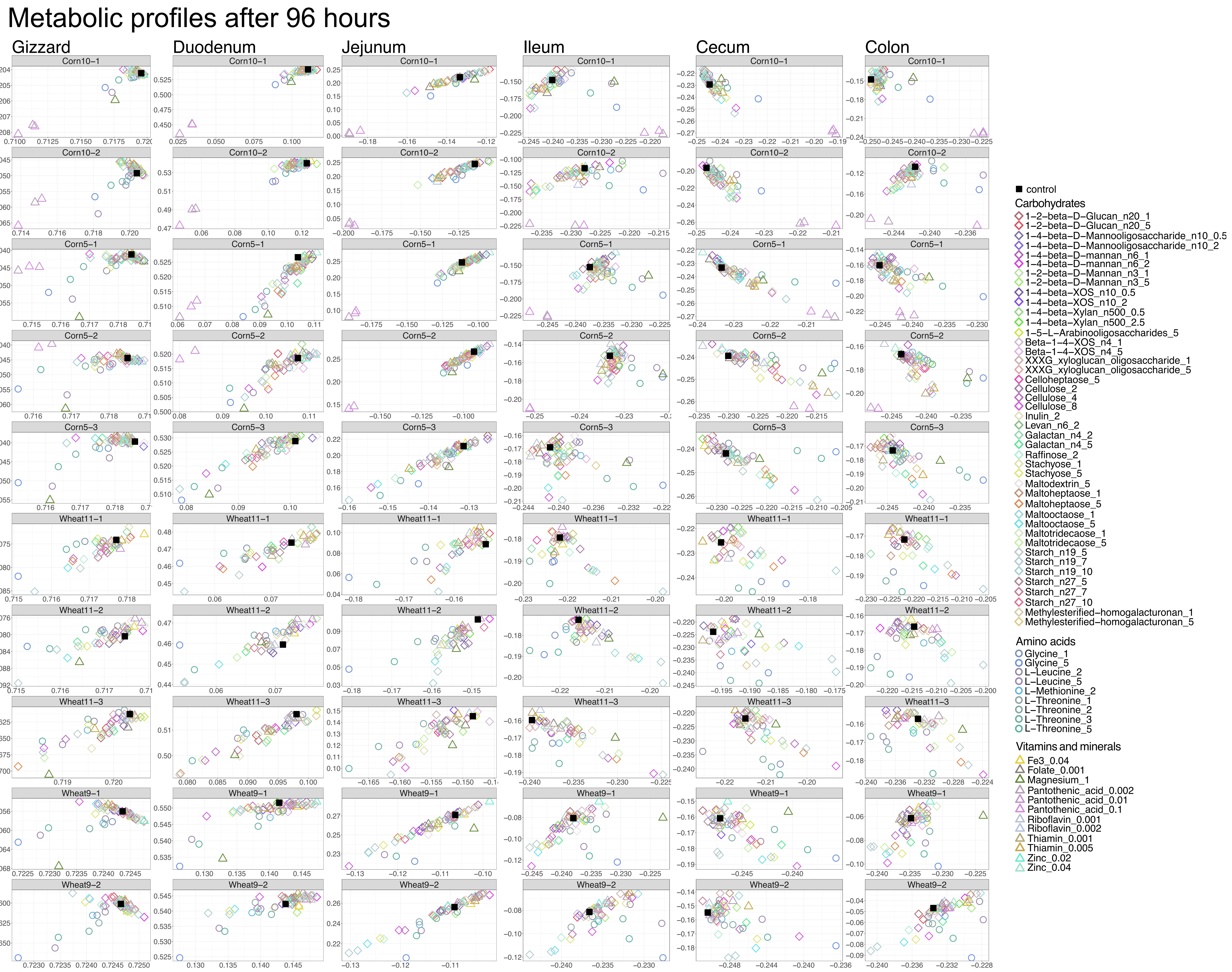

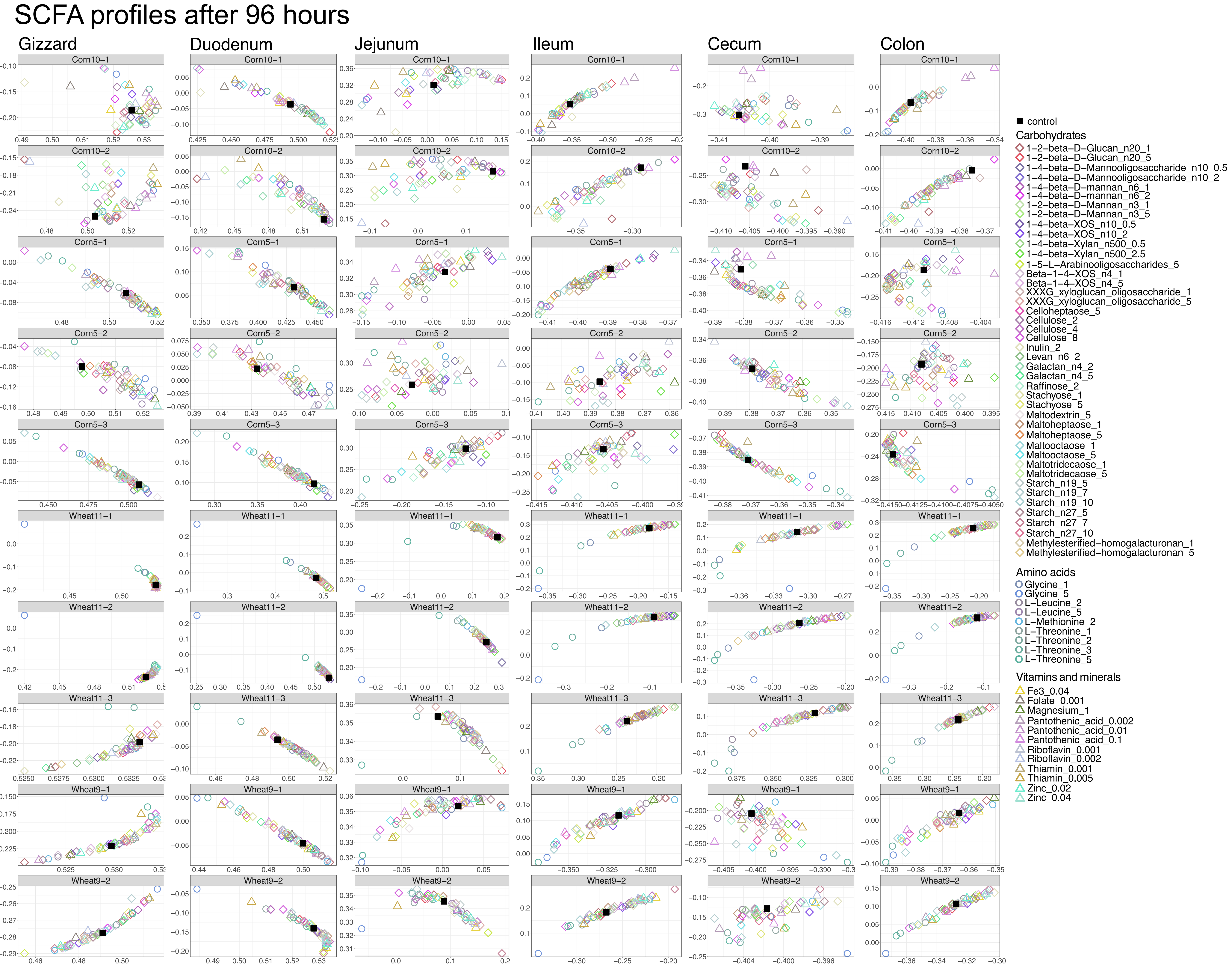

### Supplemental Figure 5

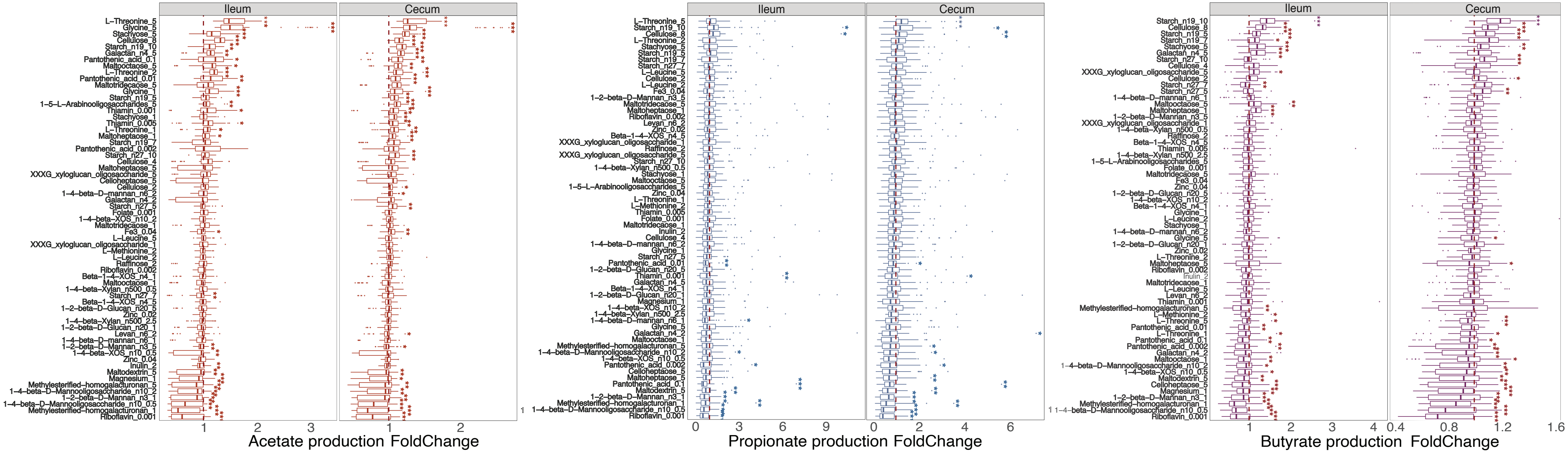

### Supplemental Figure 6

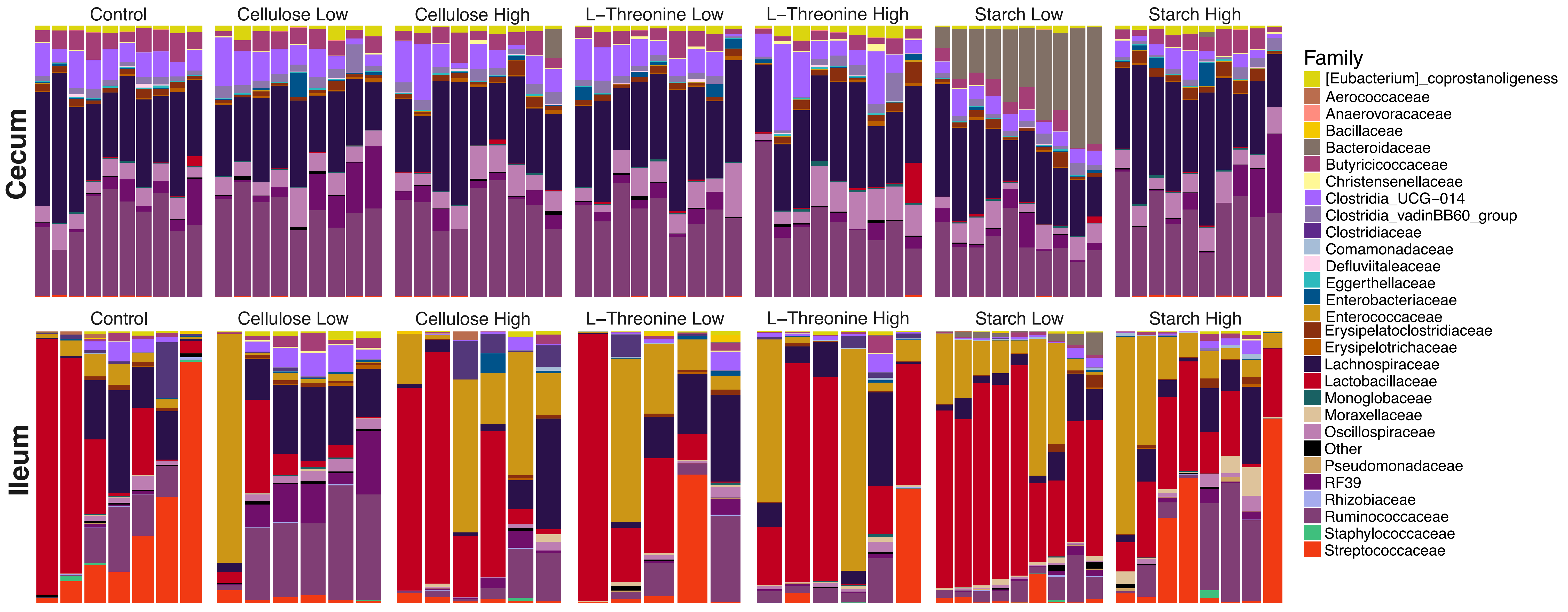

### Supplemental Figure 7

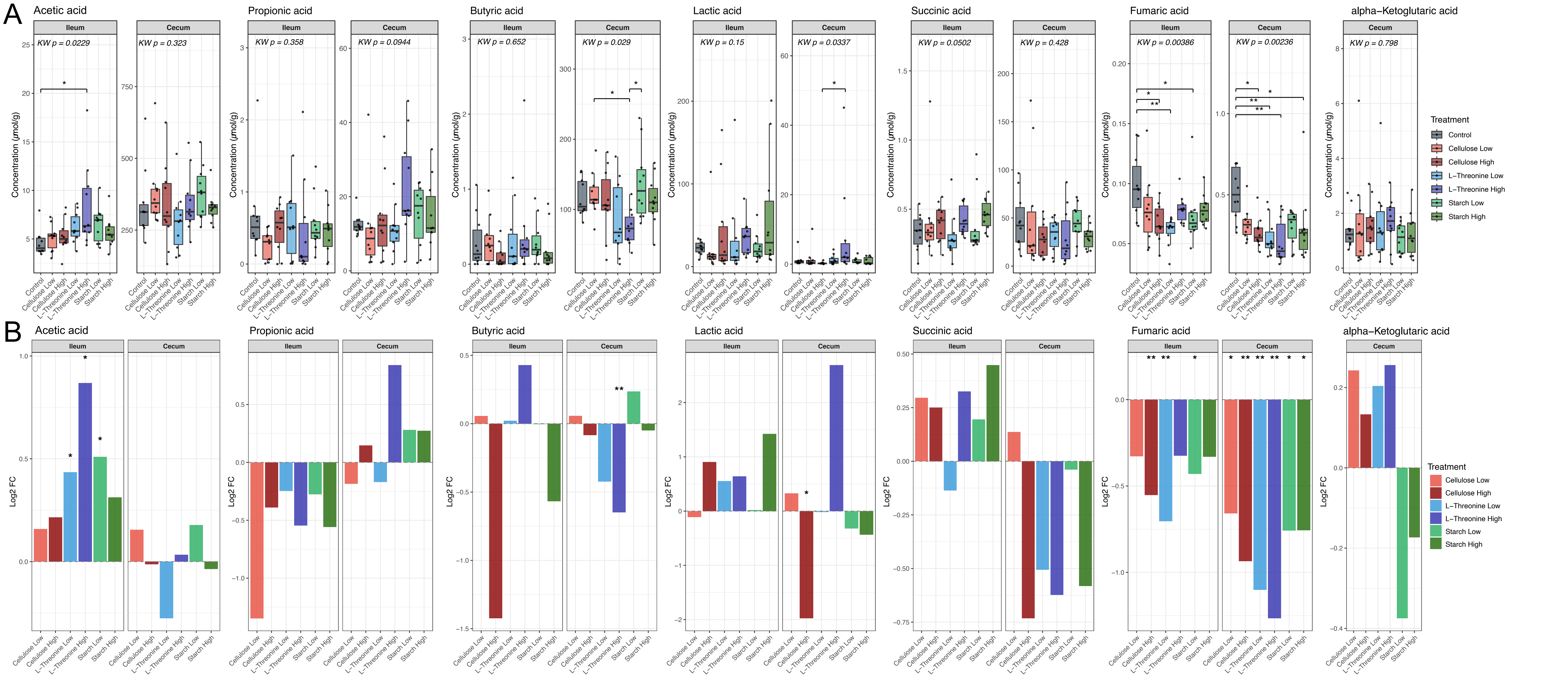

### Supplemental Figure 8

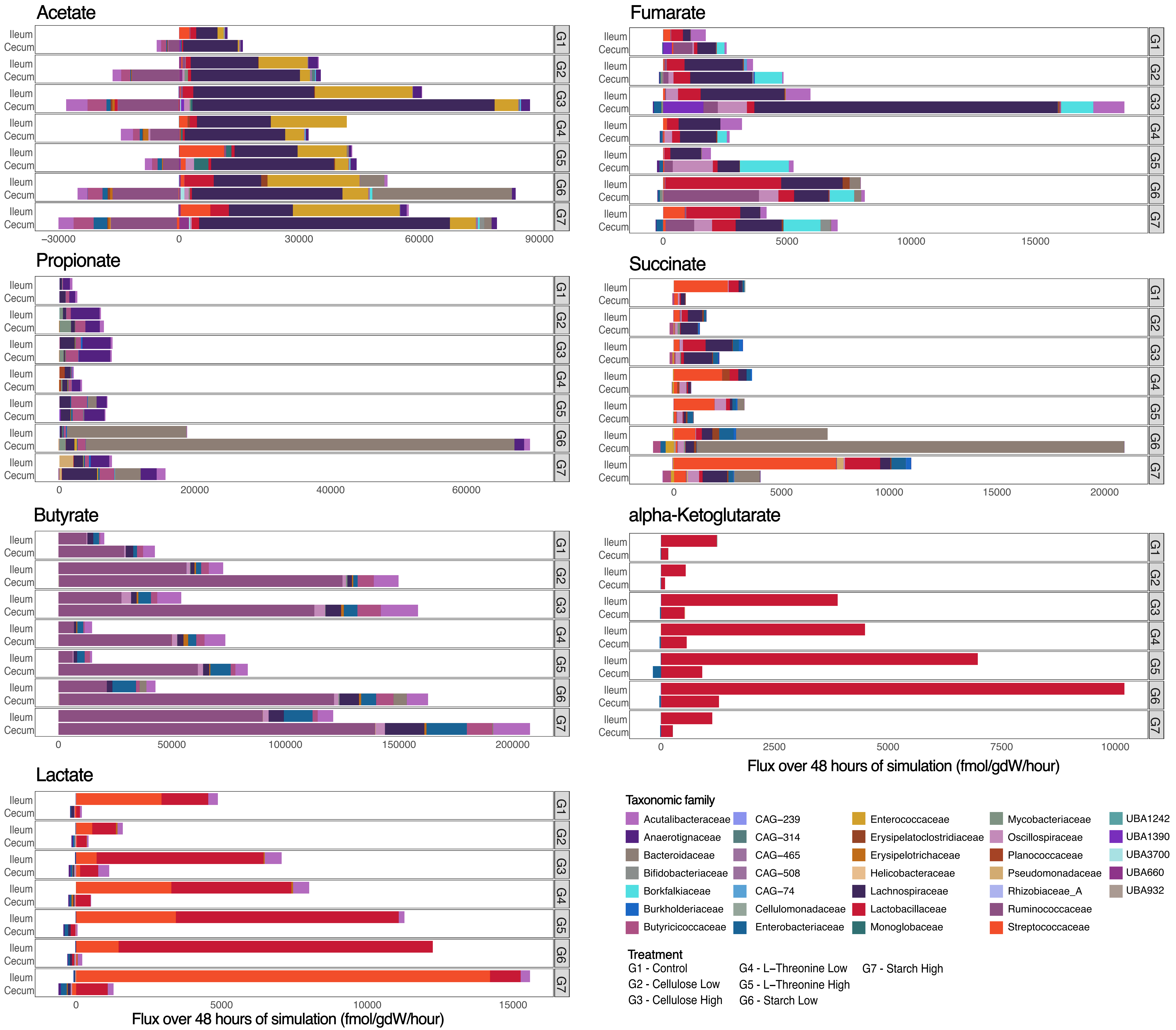
